# Life-history predicts global population responses to the weather in the terrestrial mammals

**DOI:** 10.1101/2021.04.22.440896

**Authors:** John Jackson, Christie Le Coeur, Owen R Jones

## Abstract

With the looming threat of abrupt ecological disruption due to a changing climate, predicting which species are most vulnerable to environmental change is critical. The life-history of a species is an evolved response to its environmental context, and therefore a promising candidate for explaining differences in climate-change responses. However, we urgently need broad empirical assessments from across the worlds ecosystems to explore these predictions. Here, we use long-term abundance records from 157 species of terrestrial mammal and a two-step Bayesian meta-regression framework to investigate the link between annual weather anomalies, population growth rates, and species-level life-history. Overall, we found no consistent effect of temperature or precipitation anomalies on annual population growth rates. Furthermore, population responses to weather anomalies were not predicted by phylogenetic covariance, and instead there was variability in weather responses for populations within a species. Crucially, however, long-lived mammals with smaller litter sizes had responses with a reduced absolute magnitude compared to their shorter-living counterparts with larger litters. These results highlight the role of species-level life-history in driving responses to the environment.

## Introduction

Climate change is one of the greatest challenges we face in the twenty first century (Díaz et al., 2019). Although habitat loss and direct exploitation are currently the greatest drivers of extinction in the natural world (Daskalova, Myers-Smith, Bjorkman, et al., 2020; Díaz et al., 2019), changes to the climate are predicted to cause widespread declines to global biodiversity in the coming decades (Almond et al., 2020; Soroye et al., 2020; Thomas et al., 2004). For mammals and birds, temperature increases are already associated with declining population trends (Spooner et al., 2018) and many endangered species have already been negatively impacted by climate change in at least part of their range (Pacifici et al., 2017). Perhaps more worryingly, abrupt ecological disruption due to runaway climate change has been predicted to have large negative impacts on biodiversity, with tropical ecosystems being affected as early as 2030 (Trisos et al., 2020). Furthermore, these future impacts will likely be exacerbated by synergism between the climate and other drivers of extinction such as habitat loss (Brook et al., 2008; Williams et al., 2020). Research highlighting the species and ecosystems that are most vulnerable to climate change impacts will therefore provide crucial knowledge to prevent future losses to global biodiversity.

Not all species are equally vulnerable to climate change. Species vary in their climatic niches and in their behavioural, physiological, and demographic responses to environmental change and we therefore expect there to be both climate ‘winners’ and ‘losers’ (Antão et al., 2020; Bellard et al., 2012; Moritz & Agudo, 2013). At a macro-scale, species occupancy data highlight that geographic range shifts are the key response associated with climate change across taxa, resulting in changes to community composition, but not necessarily population decline (Chen et al., 2011; Dornelas et al., 2019). Recent broad assessments of biodiversity change metrics from species assemblage and abundance records mirror this paradigm with both occupancy and abundance trend patterns suggesting a balanced frequency of winners and losers (Dornelas et al., 2019; Leung et al., 2020). In the Marine realm, species richness increases were associated with warming temperatures, consistent with the expectation that warming marine ecosystems will receive an influx of species tracking suitable temperatures (Antão et al., 2020). However, there was no consistent temperature-related biodiversity change effect on land (Antão et al., 2020). Changes in species richness and other biodiversity metrics however do not necessarily equate to population declines. There was evidence that weather covariates improved the prediction accuracy of time-series abundance models for 492 animal time-series, but there was a high risk of overfitting in short time-series (Knape & de Valpine, 2011). For birds and mammals average abundance trends were negatively associated with rates of climate warming (Spooner et al., 2018). Studies unpicking how changes in weather patterns cause population change are therefore vital (Coulson et al., 2001), and a growing body of literature is exploring the relationship between the climate and the demographic processes driving population decline (Cordes et al., 2020; Layton-Matthews et al., 2020; Paniw et al., 2019, 2021; Woodroffe et al., 2017). Applying these concepts at a comparative scale and assessing finer-scale population changes with respect to changes in the weather, and their relationship to species traits, will aid in illuminating consistent or disparate climate change responses across the tree of life (Compagnoni et al., 2021; Paniw et al., 2021).

Variation in the demographic responses of organisms climate change suggests that life-history is a promising target for explaining responses to environmental change. The timing of demographic events relating to the key demographic rates of survival and recruitment are evolved responses to the environment, and characteristics relating to both ‘slow’ and ‘fast’ life-histories are therefore adaptive in different environmental contexts (Stearns, 1992). Indeed, Pacifici et al.(2017) concluded that intrinsic traits, including habitat specialisation and aspects of life-history, were associated with negative climate-mediated population effects reported in mammals and birds. Life-history differences between three amphibian species in Western Europe drove differences in survival and reproduction in response to the North Atlantic Oscillation (Cayuela et al., 2017). Generally, we expect that organisms with slower life-histories are better-adapted to cope with fluctuations in the environment. Longer-lived organisms have a reduced relative effect of variability in vital rates, variability which is expected during environmental change, on population growth rates (Morris et al., 2008) and long-lived plants have weaker absolute demographic responses to weather (Compagnoni et al., 2021). However, while generally buffered, long-lived, slow reproducing animals are often more at risk of extinction (Cardillo et al., 2000), and slower to recover when perturbed (Gamelon et al., 2014; Jackson et al., 2019; Turkalo et al., 2016). Furthermore, the recently developed concept of demographic resilience uses demographic rates characterising the life cycle of an organism to quantify their resilience to perturbations (Capdevila et al., 2020). Comparative approaches linking life-history traits to climate change responses may therefore provide a crucial predictive link to improve our understanding of climate vulnerability.

In this study, we investigated annual population responses to temperature and precipitation in populations of terrestrial mammals across the world’s ecosystems. Importantly, we tested whether life-history predicts population responses to the weather, and therefore its utility in assessing vulnerability to climate change. We addressed these questions using 486 long-term (≥10 years) abundance records from 157 species of terrestrial mammal obtained from the Living Planet Database (Almond et al., 2020), by implementing a two-step meta-regression framework. First, for each abundance record, we assessed how observed annual population growth rates were influenced by weather anomalies (annual deviation from long-term average weather patterns) using autoregressive additive models that accounted for temporal autocorrelation in abundance records and overall abundance trends. Then, we used a phylogenetically controlled Bayesian meta-regression with weather effect coefficients as the response variable to address three key questions: 1) Are there consistent temperature and precipitation effects on abundance change across the terrestrial mammals? 2) How are these patterns influenced by covariance both within and between species, and are there vulnerable biomes? 3) Can species-level life-history traits predict the magnitude of population responses to the weather? The terrestrial mammals are an ideal study system to explore the predictors of population responses to climate change because they are a well-studied group with a combination of intensive abundance monitoring across the globe (Almond et al., 2020), detailed life-history information for hundreds of species (Conde et al., 2019; Myhrvold et al., 2015) and a highly resolved phylogeny to facilitate phylogenetic comparative analyses (Upham et al., 2019). Furthermore, there is growing evidence from the mammals of the mechanistic links between the climate, demography, and population dynamics (Coulson et al., 2001; Paniw et al., 2019, 2021; Woodroffe et al., 2017).

## Results

We assessed population responses to weather in 486 long-term abundance time-series records of 157 species of terrestrial mammals from across the world’s ecosystems (Fig. 1). The time-series records ranged in duration from 10 years to 35 years, with mean and median record lengths across records of 15.7 and 14 years, respectively (Fig. 1). The records were distributed across 13 terrestrial biomes, including both tropical and temperate regions, but were generally biased towards north western Europe and North America. We had records from 12 of 27 mammalian orders recognised by the IUCN Red List for threatened species (IUCN, 2016), but most densely in the Artiodactyla (n = 172), Carnivora (n = 127) and Rodentia (n = 82) (Fig. 1). The number of records for each species ranged from 1-17, with a mean of 3.1 and median of 2 records per species.

**Figure 1.**
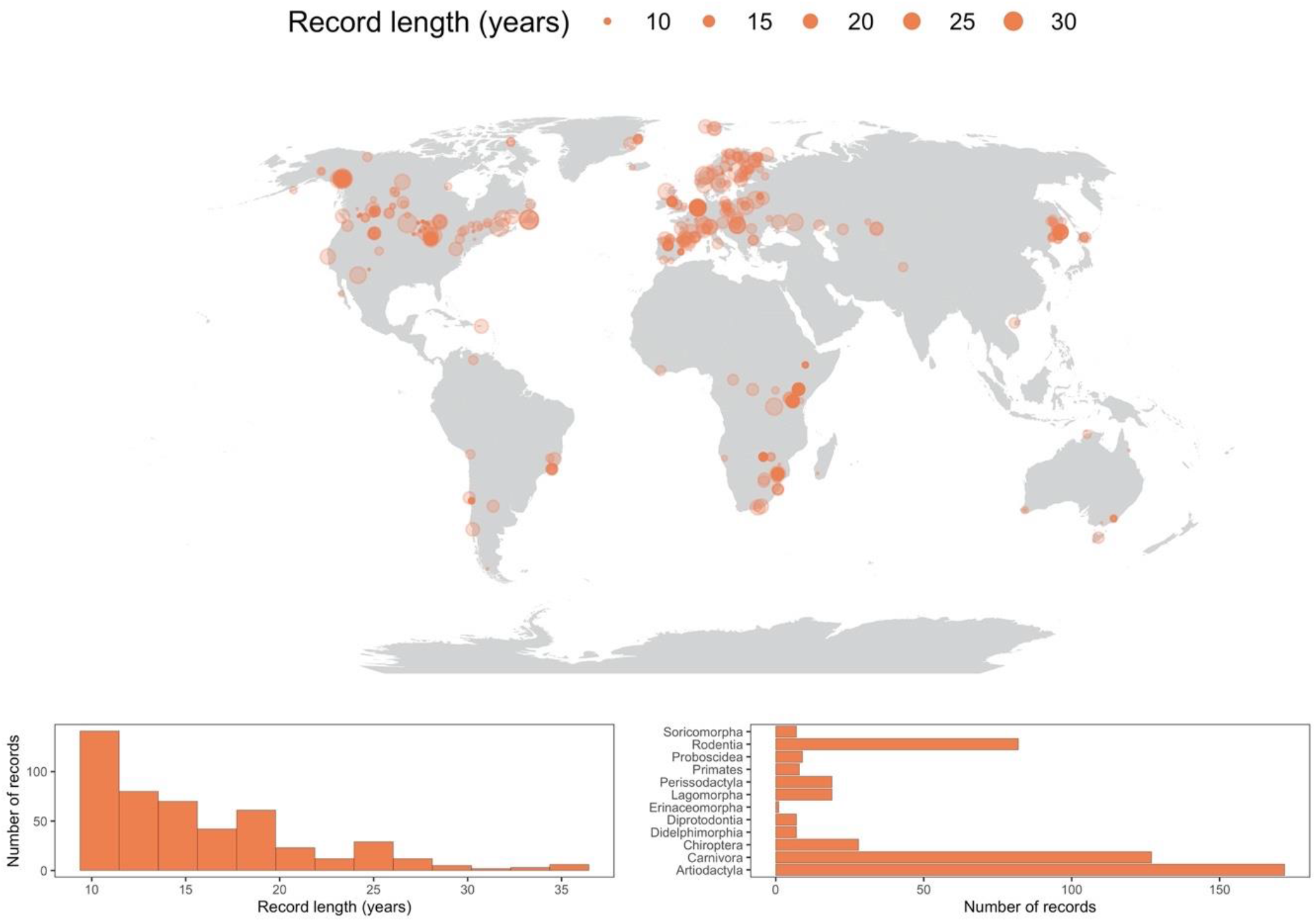
486 long-term abundance records for the terrestrial mammals. Map gives the coordinate locations for each record analysed in the current study. The size of the point gives the record duration in years. The histogram in the bottom left gives the distribution of record lengths across the whole dataset. The bar graph in the bottom right is a frequency distribution of each of the mammal orders analysed in the current study.

### No consistent population response to weather

Overall, we found no consistent effect of either temperature or precipitation anomalies on annual population growth rates in the terrestrial mammals (Fig.2). The raw weather effects on population growth rates, *ω*, varied across species and records but were centred close to 0, with a mean temperature coefficient of −0.32 (±1.76 SD) and mean precipitation coefficient of 0.07 (±0.81 SD). Furthermore, 95% of records had temperature and precipitation coefficients between −4.29-3.17 and −1.41-1.88, respectively. Nevertheless, approximately 8% (n = 42) of temperature effects and 1% of precipitation were greater than 3 or less than −3, indicating that small clusters of populations experienced more extreme annual responses to the weather (Fig.2). Our Bayesian meta-regression, controlling for both within species variance, phylogenetic covariance and differences in sample size (number of years) between records, mirrored the lack of consistent weather effects on population growth. The posterior mean global intercept, 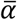, for temperature effects was 0.02 [−0.21-0.25] (95% credible intervals) and for precipitation effects was −0.07 [−0.31-0.15] (Fig.2a and Fig.2b). There was, however, a positive effect of the number of years of population data for a record and the response to temperature, with a linear slope, *β*_*N*_ of 0.12 [0.03-0.21]. Together with the results of the global intercept 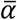, this suggests that shorter records were associated with more negative temperature effects. Overall, these results highlight the paradigm of the existence of both winners and losers in weather responses, but no clear effect across the Mammalia.

**Figure 2.**
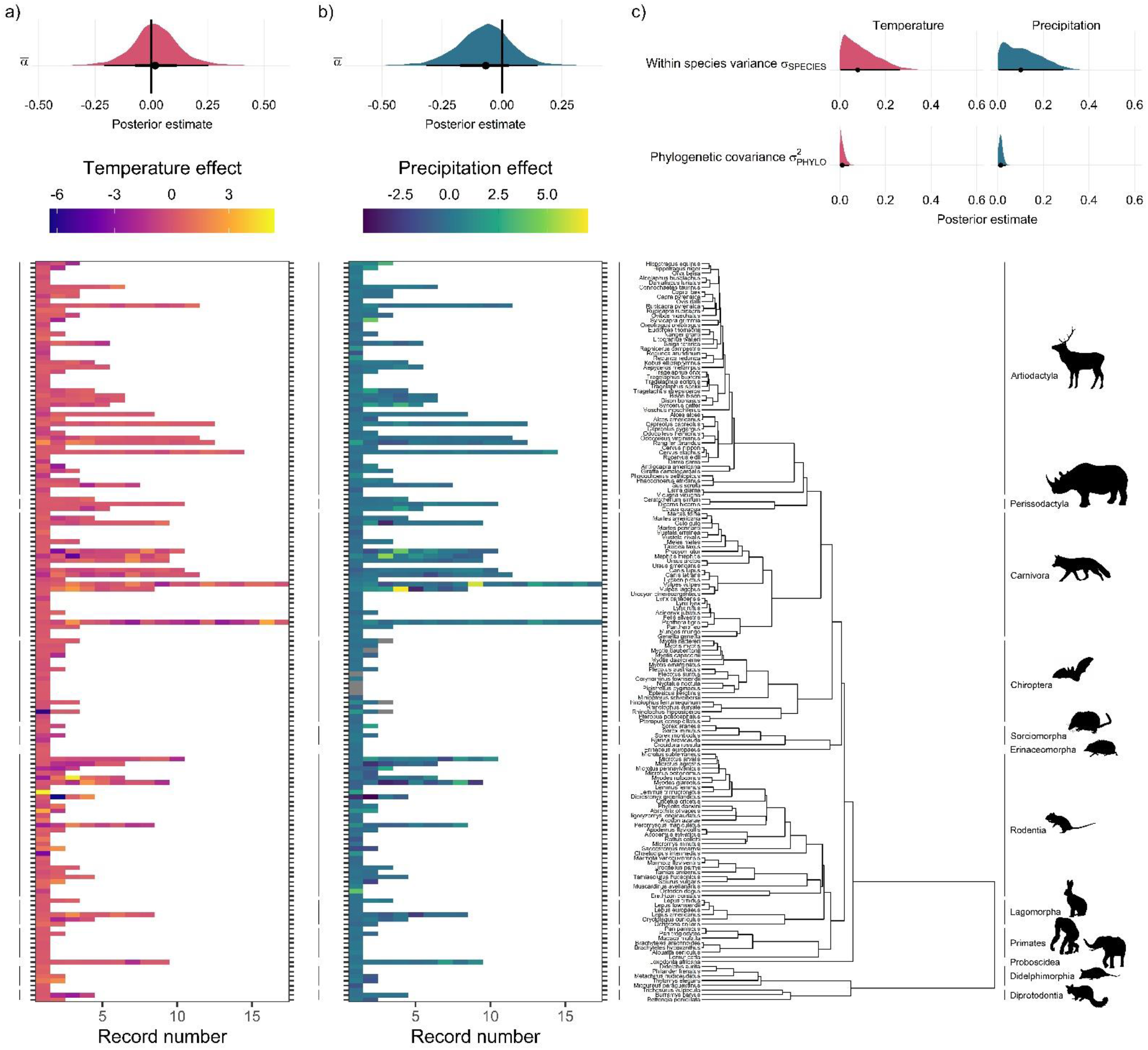
Global population responses to weather in the terrestrial mammals. Heatmaps for population responses to temperature (a) and precipitation (b) for 157 species of terrestrial mammal. Each row of the heatmap is a species, and coloured rectangles are the population records. The colour denotes the coefficient of temperature/precipitation effects derived from autoregressive additive models. Here, positive numbers indicate that positive temperature/precipitation anomalies (hotter/wetter than average in a given year) were associated with increases in population size, and *vice versa*. The distribution half-eye plots in (a) and (b) (top) are summaries of the posterior distribution for the global intercept (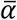) of temperature and precipitation responses across records, fit with a Gaussian Bayesian meta-regression. The points give the approximate posterior mean and the error bar is calculated using a cumulative distribution function. Bayesian models were fit incorporating phylogenetic covariance using the maximum clade credibility tree from Upham et al.(2019), which is plotted on the right with annotations indicating the mammal order. The distribution half-eye plots in (c) are the posterior distribution summaries for phylogenetic covariance and within-species variance from the Gaussian Bayesian meta-regression.

### Spatial effects and variation between species

We tested whether there were differences in weather responses among ecological biomes because biome effects may be indicative of more extreme responses to weather in some habitats. Using leave-one-out cross-validation, we compared the predictive performance of the model including the effect of biome relative to the base model, and we found no evidence for an influence of biome on either temperature (Δelpd = −0.67 relative to base model) or precipitation (Δelpd = −0.73) effects (see Fig. S16-17 for more information). Furthermore, we explored the role of spatial autocorrelation at driving differences in weather coefficients across records using Morans I tests and spatially explicit meta-regressions but did not find evidence for spatial autocorrelation in weather effects (Figs. S19-S21). We also incorporated both phylogenetic covariance 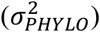 and species-level variance (*σ*_*SPECIES*_) to capture both among and within-species variation. Interestingly, we found far greater levels of within-species variation in temperature responses compared to among-species variance (Fig. 2c). The posterior mean for species-level variance in temperature effects was 0.2 [0.01-0.4] which was 20 times greater than the posterior estimate of 0.01 [0.0-0.03] for phylogenetic covariance (Fig. 2c). Similarly, for precipitation the posterior mean for species-level variance was five times greater than for phylogenetic covariance, with a value 0.05 [0.0-0.15] compared to 0.01 [0.0-0.02] (Fig. 2c). These patterns are reflected in the temperature and precipitation coefficients, for which large variation can be seen among records of the same species. For example, *Myodes glareolus* (bank vole) in the Rodentia had 9 population records, and a range of temperature/precipitation effects of −3.33-3.86 and −2.72-2.41 respectively, compared to −11.60-9.22 and −3.47-3.22 across Rodentia as a whole (Fig.2). This result highlights the potential importance of within-species variability in population responses to environmental change.

### Life-history predicts population responses to weather

Across the terrestrial mammals, we found that both maximum longevity and mean litter size at the species-level predicted the magnitude of population responses to weather. We tested a set of Gamma models incorporating univariate, multivariate and 2-way interaction effects of maximum longevity, litter size, and adult body mass and their influence on the absolute magnitude of temperature/precipitation effects using model selection and leave-one-out cross-validation (Table S1 & S2). As with our Gaussian models of overall weather effects, we found that the sample size of a record had strong negative impact on the absolute temperature and precipitation responses, with posterior estimates on the linear predictor scale of *β*_*N*_ = −0.30 [−0.38- −0.21] and *β*_*N*_ = −0.37 [−0.47- −0.26], respectively (Fig. S18). Namely, shorter records were associated with temperature and precipitation responses of a larger magnitude. We found no association between adult body mass and either temperature (*β*_*BODYMASS*_ = −0.02 [−0.15-0.10]) or precipitation responses (*β*_*BODYMASS*_ = −0.00 [−0.17-0.17]). Furthermore, we found no strong evidence for any two-way interactions between life-history variables (Table S1 & S2). For both temperature and precipitation effects, the most competitive model was the univariate model including maximum longevity (Δelpd = 5.44 and Δelpd = 1.03, compared to the base model for temperature and precipitation, respectively; Table S1 & S1). However, univariate models including litter size also had a higher predictive performance than the base model (Δelpd = 3.98 and Δelpd = 0.8 for temperature and precipitation, respectively). For temperature, the second-best predictive model was the one that included univariate effects for longevity, bodymass and litter size (Δelpd = 4.54; Table S1), and this model was also competitive for precipitation (Δelpd = 0.69; Table S2). Therefore, in both cases we selected the models including all univariate life-history effects.

For both temperature and precipitation, our results highlight that shorter-living mammals with greater litter sizes experienced weather effects of a greater magnitude than longer-living, slowly reproducing mammals (Fig. 3). The magnitude of weather responses was negatively associated with longevity, with posterior means on the linear predictor scale of *β*_*LONGEVITY*_ = −0.20 [−0.41-0.02] and *β*_*LONGEVITY*_ = −0.17 [−0.42-0.09] for temperature and precipitation, respectively (Fig. 3a & 3c). Thus, a maximum longevity change from 10 months (*Akodon azarae*) to 80 years (*Loxodonta africana*) was associated with a 2.36-fold and 2.05-fold decrease in the predicted magnitude of responses to temperature and precipitation. So, for every additional 5 years of life, there was a 16.8% decrease in the magnitude of responses to temperature and 14.6% decrease in the magnitude of responses to precipitation. An organism’s longevity is strongly correlated to their body mass, but the effect of longevity held irrespective of whether adult body mass was also included in the model. In contrast, but also following key predictions from life-history theory, the magnitude of weather responses had a positive association with litter size, with posterior means of *β*_*LITTER*_ = 0.16 [0.02-0.32] and *β*_*LITTER*_ = 0.11 [−0.08-0.30] for temperature and precipitation, respectively (Fig. 3b & 3d). In other words, mammals bearing more offspring in a single litter had greater responses to temperature and precipitation. A change in litter size from 1 (monotocous species, various) to 17 (*Thylamys elegans*) was associated with a 1.99-fold and 1.60-fold increase in the predicted magnitude of temperature and precipitation responses. For every additional offspring invested into at the litter stage, there is a 12.4% increase in the magnitude of temperature responses and 10% increase in the magnitude of precipitation responses.

**Figure 3.**
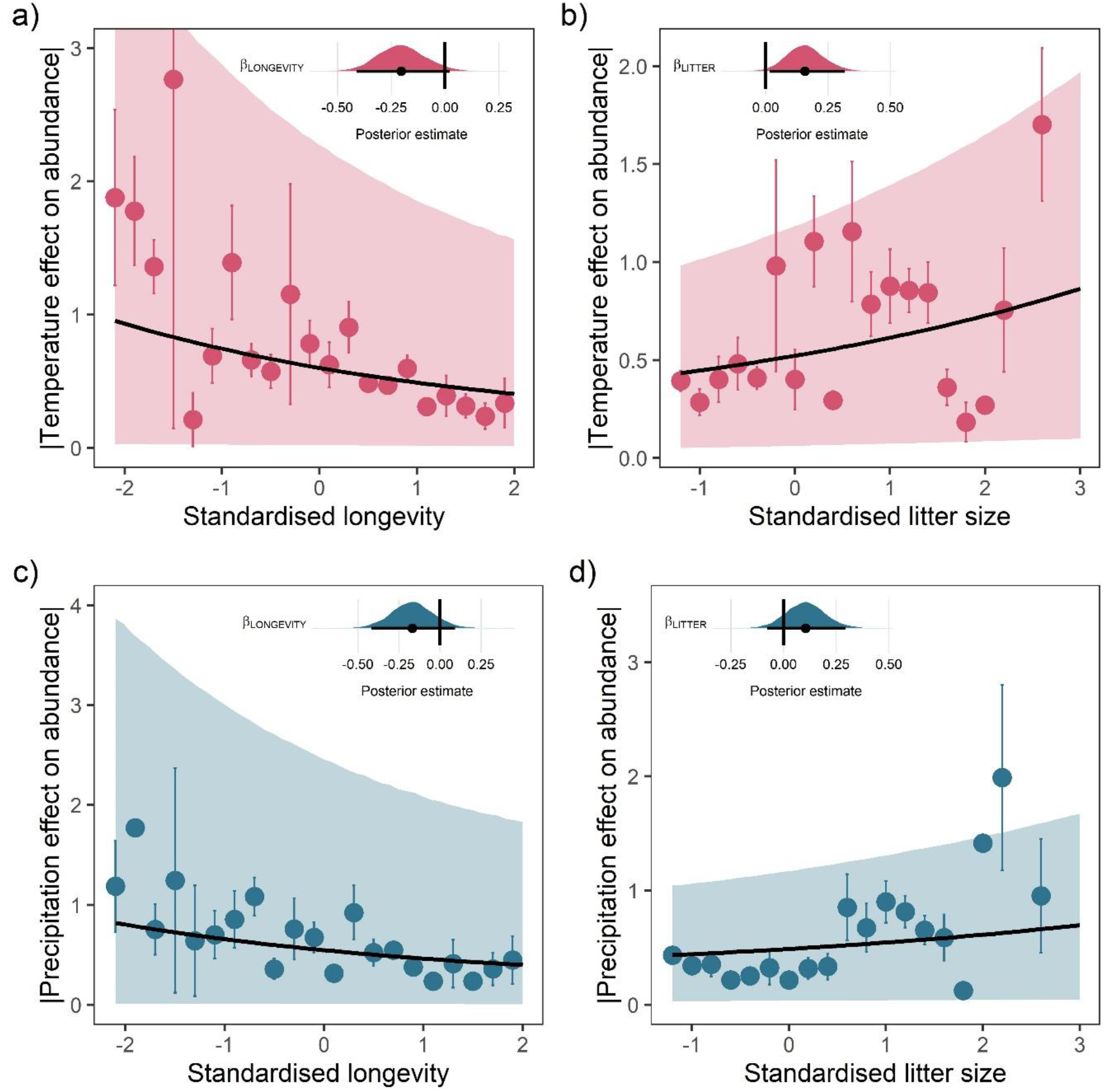
Life-history predicts population responses to weather in the terrestrial mammals. Each panel presents the mean absolute effect of temperature (a & b) and precipitation on population growth rates, │*ω*│, for standardised maximum longevity (a & c) and standardised mean litter size (b &d). Standardisation was performed using a z-transformation of natural-log of the raw life-history trait. The values on each x-axis are split into equal bins of 0.2 from the minimum to the maximum life-history value. Points are coefficient means, with standard error bars. The black lines are the mean posterior predictions from the best predictive model, where predictions were calculated averaging over all other covariates and varying effects in the model. The shaded intervals are the 80% quantile prediction intervals. Panel insets give posterior distribution summaries for the slope terms presented in each panel. Two points are omitted from the plotting panel due to large mean coefficient values and high standard errors, which are visible on the plot.

## Discussion

Our results provide an important empirical link between a species’ life-history and its population responses to environmental change. While we found no consistent patterns of responses to temperature and precipitation anomalies across the mammals, life-history traits relating to the pace of life were associated with responses to weather. Namely, shorter-living species with increased litters sizes, or species characterised with ‘fast’ life-history traits, responded with a greater magnitude compared to those with ‘slow’ life-history traits. While it has long been theorised that an organism’s life-history traits evolve in response to, and as an adaptation to, environmental conditions (Stearns, 1992), rarely has this theory been tested at a global scale. We find strong support for the hypothesis that longevity, and ‘slow’ life-history characteristics more generally, buffer organisms against short-term variability in the environment (Morris et al., 2008) and add to a small number of studies linking population demography and the climate (Compagnoni et al., 2021; Paniw et al., 2021). We predict that in the short term abrupt ecological disruption from climate change will have a disproportionate impact the abundance of shorter-lived species with higher reproductive output. We do not argue that long-lived species are less vulnerable to climate change. Over longer time-scales, species with slow life-history are also slower to recover from perturbations (Gamelon et al., 2014). However, critically our results highlight the potential utility of life-history traits for predicting species vulnerability to climate change.

Demography has a vital role to play in predicting population declines in the Anthropocene and in highlighting targets for conservation management (Conde et al., 2019; Richards et al., 2021). Our study emphasises this role, demonstrating the predictive power of life-history traits when investigating responses to environmental change. However, there are limitations and barriers to the utility of demography in conservation. Only 1.3% of tetrapods globally have sufficient demographic information with which to estimate population dynamics (Conde et al., 2019). Here, we used summary traits that are available for many species (maximum longevity and mean litter size), but in particular maximum recorded longevity, while sufficient as a broad indicator, is strongly influenced by sampling variance and a flawed measure of longevity differences between taxa (Moorad et al., 2012). Ideally, lifetables with mortality and reproduction trajectories across the lifecycle can be combined with data on external drivers to investigate detailed patterns in population dynamics, rather than relying on abundance trends (Desforges et al., 2018; Jackson et al., 2019). The recent development of the demographic resilience framework, which uses demographic data across the lifecycle to simulate how a population may respond to perturbations (Capdevila et al., 2020), has excellent potential in extending these findings to explore demographic relationships with climate responses in detail. Unfortunately, however, detailed (st)age-specific demographic information is not currently available for a majority of species, but growing in availability rapidly (Salguero-Gómez et al., 2016). Therefore, there is a need to continue to increase the collection of demographic data (and other traits) for many more species than are currently available (Conde et al., 2019), so that we may predict population changes with respect to environmental change. Achieving this target may revolutionise the way we quantify species vulnerability to climate change (Antão et al., 2020; Dornelas et al., 2019; Leung et al., 2020; Paniw et al., 2021), helping to prevent extinctions before they occur.

In line with recent global assessments of biodiversity in the face of climatic change (Paniw et al., 2021), we did not find an overall consistent effect of weather anomalies on population growth rates. This may in part reflect the fact that abundance changes are a higher-order process determined by complex interactions between demographic processes that counteract each other (Leung et al., 2020; Paniw et al., 2021). However, our results contrast with findings of linear associations between mammal abundance and temperature change (Spooner et al., 2018). These differences may reflect our approach to investigate annual changes, rather than long-term trends. Significant population trends from long time-series are detectable from smaller component time-series even when sampling is incomplete (Wauchope et al., 2019), and thus responses detected in trends may reflect broader changes in response to the climate that are not detected in models of annual change. Furthermore, we estimated linear, annual effects of weather on population growth rates, where population responses may actually be more complex non-linear patterns or lagged effects. However, the detection of climate effects on average trends may also be confounded by effects of other (sometimes more dominant) drivers (e.g. habitat loss)(Daskalova, Myers-Smith, Bjorkman, et al., 2020). Nevertheless, our findings can be explained in light of recent studies from the Living Planet Database that have found that the large majority of records do not exhibit population declines (Leung et al., 2020).

Interestingly, we did not find evidence for phylogenetic covariance in weather responses between species. Recent evidence from birds indicated strong phylogenetic covariance in vital rates, particularly in adult survival, and the incorporation of phylogenetic information greatly improved predictive performance when imputing vital rates (James et al., 2020). Therefore, as with overall patterns, our findings may reflect the trade-offs between vital rates, which cancel one another out when scaling up to population-level processes such as population growth rates in response to the weather (Paniw et al., 2021). Furthermore, for long-term time-series, there may also be temporal trade-offs in vital rates, where for example investing heavily into survival in one year (in response to climate) may impact subsequent reproduction for several years, decreasing the magnitude of population growth rates. The extent of phylogenetic covariance in vital rate responses and trade-offs remains unknown, understanding how the climate impacts demographic rates across species may provide a useful tool for imputing population responses to the climate across the tree of life (James et al., 2020).

We highlight the importance of variation in population responses to climate within a species range. Sampling heterogeneity has recently been shown to have broad implications for metrics of population dynamics, where demographic rates are poorly correlated among sampling sites for the same species (Engbo et al., 2020; Römer et al., 2021). Therefore, inferences obtained from monitoring single populations or studies may not accurately portray species-level variability. This has broad implications for macroecology, particularly for population viability assessments (PVA) and species-distribution modelling. First, as well as suffering from data quality issues in their parameterisation (Chaudhary & Oli, 2020), our findings suggest that PVAs based on data from a single population may not accurately reflect population viability across a species’ geographic range. Therefore, incorporating detailed demographic data, and investigating differences in population responses across a range, could greatly improve our perspective on population viability (Desforges et al., 2018). Second, presence-only models of species distributions that do not account for the fact that responses to the environment within a species range do not accurately represent species distributions (Benito Garzón et al., 2019). Moving towards trait-based monitoring and explicitly including demographic processes with mechanistic links to appropriate drivers into species distribution models could greatly improve predictions of climate change impacts on the biosphere (Trisos et al., 2020).

Ultimately, improving our predictions of how humans are influencing the natural world is paramount to prevent rapid declines to global biodiversity (Kissling et al., 2018). This however requires a large shift towards both broad and detailed monitoring of global biodiversity. We show that linking species traits such as life-history to changes in the environment may equip us with tools to predict and prevent future losses.

## Materials and Methods

To assess the effects of weather on population growth rates we collated information on global weather and the abundance, life-history and phylogeny of the terrestrial mammals. All analyses were carried out using R version 4.0.5 (R Core Team, 2021). For all data on the terrestrial mammals, taxonomies were resolved using the *taxize* package version 0.9.98 (Chamberlain et al., 2020) and matched using the Global Biodiversity Information Facility database (https://www.gbif.org/). All code used in the current study and full descriptions of the analyses are archived in the Zenodo repository (doi:10.5281/zenodo.4707232), which was created from the following GitHub repository https://github.com/jjackson-eco/mammal_weather_lifehistory.

### Data selection

For full descriptions of the data selection process please refer to S1. Long-term annual time-series abundance data from across the terrestrial mammals were obtained from the Living Planet Database found at https://livingplanetindex.org/data_portal. Abundance is measured in several ways (e.g. population counts and density, which does not impact population responses, see Fig. S19), and so we natural-log-transformed population growth rates to ensure that weather effects were comparable across records. Our final dataset contained 486 geo-referenced records from 157 terrestrial mammal species, which was used in all subsequent analyses (Fig. 1).

Global weather data was obtained from version 1.2.1 of the CHELSA monthly gridded temperature and precipitation dataset at a spatial resolution of 30 arc seconds (~ 1km^2^) for all months between 1979-2013 across the globe’s land surface (Karger et al., 2017). Generally, we expect that organisms will respond to deviations in the weather compared to the average values, as opposed to raw weather variables. Furthermore, across the globes surface the variance in weather variables changes substantially, which may influence population responses. Thus, we explored population responses for the key weather variable of standardised annual anomalies, and then validated our approach using annual weather variance. These weather anomalies are the average deviation of the temperature and precipitation from expected values in a given year.

We used three key species-level life-history traits that are available for a large number of species: maximum longevity, mean litter size and mean adult body mass. Life-history data were collected from Conde et al. (2019), Myhrvold et al. (2015) Jones et al. (2009) and Tacutu et al. (2013). For analyses we z-transformed the natural-logarithm of raw life-history trait data. The mammal phylogeny used was the maximum clade credibility tree from Upham et al. (2019).

### Weather effects on annual population growth rates

To assess comparative population responses to weather in the terrestrial mammals we used a two-step meta-regression approach. First, for each record we estimated the effect of annual weather anomalies (and weather variance) on population growth rates. We calculated the standardised proportional population growth rate *r* in year *t* as

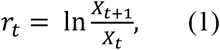

where *X* is the abundance in year *t*, transformed to prevent observations of 0.

Then, with *r*_*t*_ as the response variable, we estimated the effect of temperature and precipitation anomalies on population growth using generalised additive mixed models (GAMMs) fit using the *gamm* function of the *mgcv* package (Wood, 2017). We opted to use a general linear-modelling framework as opposed to a state-space approach, which is often employed for time-series to account for measurement error and estimate trends (see Daskalova, Myers-Smith, & Godlee, 2020). The primary reason for this choice was that we aimed to assess broad comparative patterns in population change, and did not expect systematic errors in model parameters due to measurement error. Furthermore, Daskalova, Myers-Smith, & Godlee (2020) found that abundance trend terms were highly correlated between linear and state-space approaches across the LPD, which would be expected if there are not systematic errors in measurement across the database. We did however test the implication of this choice by employing a state-space approach (see S2).

Changes in abundance are influenced by several drivers of population dynamics including habitat loss (Daskalova, Myers-Smith, Bjorkman, et al., 2020) and population processes such as density dependence (Brook & Bradshaw, 2006), which may confound any influence of the weather on abundance. Therefore, because we aimed to assess the isolated impact of weather anomalies, accounting for these trends in abundance and temporal autocorrelation was crucial. We initially explored the extent of autocorrelation in abundance patterns using timeseries analysis and found evidence for lag 1 autocorrelation in abundance, but not for greater lags (Fig. S3&S4). Furthermore, we tested the potential impact of density dependence on estimating environmental effects using an autoregressive timeseries simulation and found that environmental effects were robust to density dependence even for short timeseries (Fig. S5). Thus, for each record, we model population growth rate in each year as

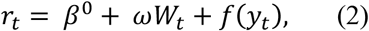

where *β*^0^ is the intercept and *ωW*_*t*_ is a linear parametric term with coefficient *ω* for the weather *W* (temperature or precipitation anomaly) in year *t*. Here, positive coefficients indicate that positive weather anomalies i.e. hotter/wetter years, were associated with population increases, and *vice versa*. Identical additive regression models were run using weather variances as the weather variable *W*. The term *f*(*y*_*t*_) captures the effect of year *y*_*t*_ as a non-linear trend, where the function *f* is a thin plate regression spline with a basis dimension of five (Wood, 2003). The function *f* was also fitted with an order 1 autoregressive (AR(1)) correlation structure, as specified in the *nlme* package (Pinheiro et al., 2014). Thus, the term *f*(*y*_*t*_) incorporates both the non-linear trend in abundance and temporal autocorrelation.

Finally, we validated our additive model approach by testing other models to calculate weather effects, including linear regressions both including and excluding temporal trends or density dependence, state-space models, and a temporally autocorrelated model fit using the *glmmTMB* package (Brooks et al., 2017) (S2; Fig. S7-S11). Weather coefficients *ω* generated using linear year effects were positively correlated to those from additive models (Fig. S9), and additive model coefficients were highly correlated with those from state-space models (Fig. S10 & S11).

### Bayesian meta-regression

Second, with the weather effects *ω* from each record as the response variable, we explored comparative patterns in population responses to weather using a Bayesian meta-regression framework implemented in the *brms* package (Bürkner, 2017). Separate models were fit for temperature and precipitation. Bayesian meta-regression was used to address three key questions: 1) Were there consistent population responses to weather across the terrestrial mammals? 2) How did population responses vary within and between species and were there spatial patterns across biomes? 3) Does life-history predict the magnitude of population responses? To address questions 1 and 2, we used Gaussian models controlling for both phylogenetic and species-level covariance. The full model for record *i* and species *j* is given by equation 3 below

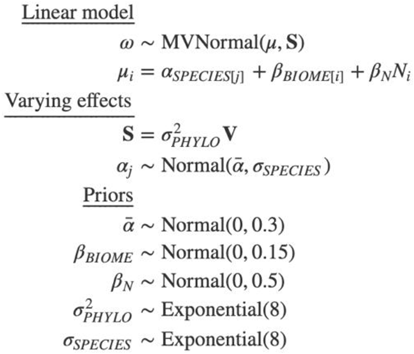

where the weather effect *ω* (z-transformed for analyses), is given by a multivariate normal distribution with mean *μ* and phylogenetic covariance matrix ***S***. The global intercept is given by 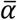, which estimates overall patterns in weather effects across records, addressing question 1. We incorporated phylogenetic covariance using a Brownian motion model, with the correlation matrix given by ***V***(calculated from the maximum clade-credibility tree) and variance factor 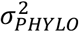, from which between-species variance was estimated. We incorporated an intercept-only varying effect for species with the term *α*_*SPECIES[j]*_, from which within-species variance was estimated with *σ*_*SPECIES*_. The term *β*_*BIOME*_ gives the spatial effect of biome on weather responses. Thus, estimating within (*σ*_*SPECIES*_) species variance, between 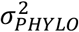 species variance and the spatial effect of biome (*β*_*BIOME*_), we explored question 2. All meta-regression models also included the linear effect of record length *N* (scaled number of years in the record) on weather effects, which was estimated using *β*_*N*_. For all meta-regression models, we used regularising priors obtained from prior predictive simulations of the slope, intercept and exponential variance terms (McElreath, 2020a, 2020b), to reflect the constraints in the raw data across species (see S3 and Fig. S12-S15 for details). Gaussian meta-regression models were also fit for weather effects calculated using the annual weather variance, and the results obtained were largely identical to those obtained for weather anomalies (Fig. S22).

For question 3, although on average we expect that species life-history influences the magnitude of responses to the environment, we have little evidence to suggest that life-history *per se* influences the directionality of responses (Morris et al., 2008). Thus, to address this question we explored how maximum longevity, litter size and adult body mass influenced the absolute magnitude of weather responses, │*ω*│, using Gamma regression models with a log link. The full model for record *i* and species *j* is given by equation 4 below

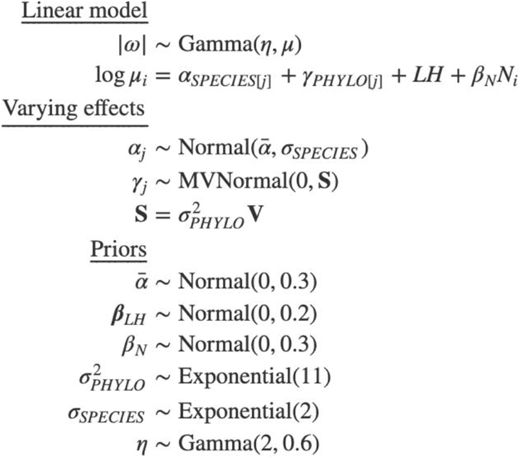

where *η* is a shape parameter that was fit with a Gamma prior, and *LH* refers to a set of linear life-history terms (*β*_1_*x*_1_ +…*β*_*k*_*x*_*k*_) that were explored using model selection. Specifically, for the three life-history traits, we explored a set of models incorporating univariate, multivariate and 2-way interaction terms, as well as a base model excluding all life-history effects. For the full set of ten candidate models please refer to the supplementary information (table S1 & S2). All life-history effects were fit with the same Normal prior, with mean 0 and standard deviation 0.3 (S2; Fig. S14). We assessed the predictive performance of candidate models using leave-one-out cross-validation implemented in the *loo* package (Vehtari et al., 2017). Models were compared using the Bayesian LOO estimate of out-of-sample predictive performance, or the expected log pointwise predictive density (elpd)(Vehtari et al., 2017). All final meta-regression models were run over 3 Markov chains, with 4000 total iterations and 2000 warmup iterations per chain. Model convergence was assessed by inspecting Markov chains, and the degree of mixing between chains using 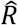.

## Supporting information

Electronic Supplementary Information

## Acknowledgements

We extend a warm thank you to the Living Planet Index team from the Zoological Society of London and the World Wildlife Foundation for the opportunity to work with this rich data source, and all the contributors to this amazing resource. We also thank Dylan Z Childs for his support on modelling the impact of density dependence on the abundance timeseries. We thank Rob Goodsell for advice on the Bayesian modelling framework, Christopher Cooney for advice on the mammal phylogeny and the Vertlife project, and Morgane Tidière for stimulating discussion and advice on the project. Thank you also to Dalia Conde and Johanna Stärk for their help with the demographic data from Conde et al. (Conde et al., 2019), and the members of the Interdisciplinary Centre for Population Dynamics (CPop), and in particular Jim Oeppen, for useful feedback on the methodology. This work was supported by the Danish Independent Research Fund.

