## Supplementary material for "Life-history predicts global population responses to the weather in the terrestrial mammals": Electronic Supplementary Information

Full analysis code and descriptions of analyses can be found archived in the following Zenodo repository (doi:10.5281/zenodo.4707232), which was created from the following GitHub repository https://github.com/jjackson-eco/mammal_weather_lifehistory.

### S1 Data selection

*Time-series abundance data*

The living planet database was developed by the World Wildlife Fund and the Zoological Society of London as a tool to monitor global biodiversity, and contains over 20,000 population records for over 4000 species of vertebrate (Almond et al., 2020). The records measure annual abundance in a variety of ways (e.g. full population counts, density, indices; these do not influence the overall effect distributions Fig. S19), and contain information on the location, realm, biome and taxonomy of the species in the record. First, we included only data for the terrestrial mammals that had species-level life-history information and coordinate locations, which referred to either specific or more general locations for each population (accounted for using weather data from a buffered radius around each location). We natural-log-transformed the raw abundance data to ensure that weather coefficients were comparable across records and abundance measure types. Because our analyses were focussed on estimating weather effects on annual population growth rates using regression models with several variables, and short timeseries are at large risk of overfitting (Knape & de Valpine, 2011) we included only long-term records with 10 or more consecutive years of abundance data, and only for years in which there was also weather data (1979-2013). In one record (for *Bettongia penicillata*), there were two blocks with ≥10 years of data, which were analysed separately. We also removed records with a high proportion (>32%) and consecutive occurrences of 0 in the raw abundance time-series. Our final dataset contained 486 records from 157 terrestrial mammal species, which was used in all subsequent analyses (Fig. S1).

*Global weather data*

The CHELSA datasets can be obtained from Karger et al.(2017). Raster files of the raw monthly mean temperature and total precipitation data were processed using the *raster, rgeos*, and *sf* packages (Bivand & Rundel, 2020; Hijmans, 2020; Pebesma, 2018). Using the Living Planet Database record coordinate locations as a centroid, we averaged the monthly weather data for a buffered radius of 5km around each record location to account for the lack of specificity in record locations using the *exactextractr* package (Baston, 2020). Averaged weather variables and weather effects for alternate buffer radii (50m and 50km) were highly correlated (Fig. S1; Fig. S6). For the anomalies, we decomposed z-transformed averaged monthly weather data for each location for the full timeseries (1979-2013) using a Seasonal-Trend Decomposition by Loess (STL). We used a seasonal window of 7 and trend window of 1000 for the decomposition and extracted the anomaly component, which describes the remainder when accounting for the trend and seasonal components of the timeseries. We then used annual mean temperature and precipitation anomalies as the key weather variables in subsequent analyses. We also investigated population responses to weather using annual weather variance, which was calculated as the annual variance of monthly mean temperature and total precipitation values.

*Species-level life-history and phylogeny*

We used three key traits that broadly characterise species-level life-history that are available for a large number of species: maximum longevity, litter size and adult body mass. We collected these traits from the compendium developed by Conde et al.(2019), combining information from three primary database sources: The Amniote Life-History Database (2015), PanTHERIA (2009) and AnAge (2013) databases. Adult body mass data was obtained exclusively from the Amniote Life-History Database (Myhrvold et al., 2015). Where multiple records were available for a single species, we took the largest maximum longevity value and the mean litter size/adult body mass. We removed erroneous raw litter size data for *Hydrochoerus hydrochaeris* and *Marmota broweri*. For analysis, we z-transformed the natural-logarithm of raw life-history trait data, and verified that the life-history variables were represented across the range of weather anomaly variables in the raw data (Fig. S2). The mammal phylogeny was obtained from Upham et al.(2019), which uses a 'backbone-and-patch' Bayesian approach for a newly assembled 31-gene supermatrix and is part of the Vertlife project (https://vertlife.org/). We used the maximum clade credibility tree in analysis, which was processed using the *ape* package (Paradis & Schliep, 2019). *Loxodonta cyclotis* (African forest elephant) was considered as *Loxodonta Africana* (African elephant) for analysis so that the abundance record and phylogenetic data matched.

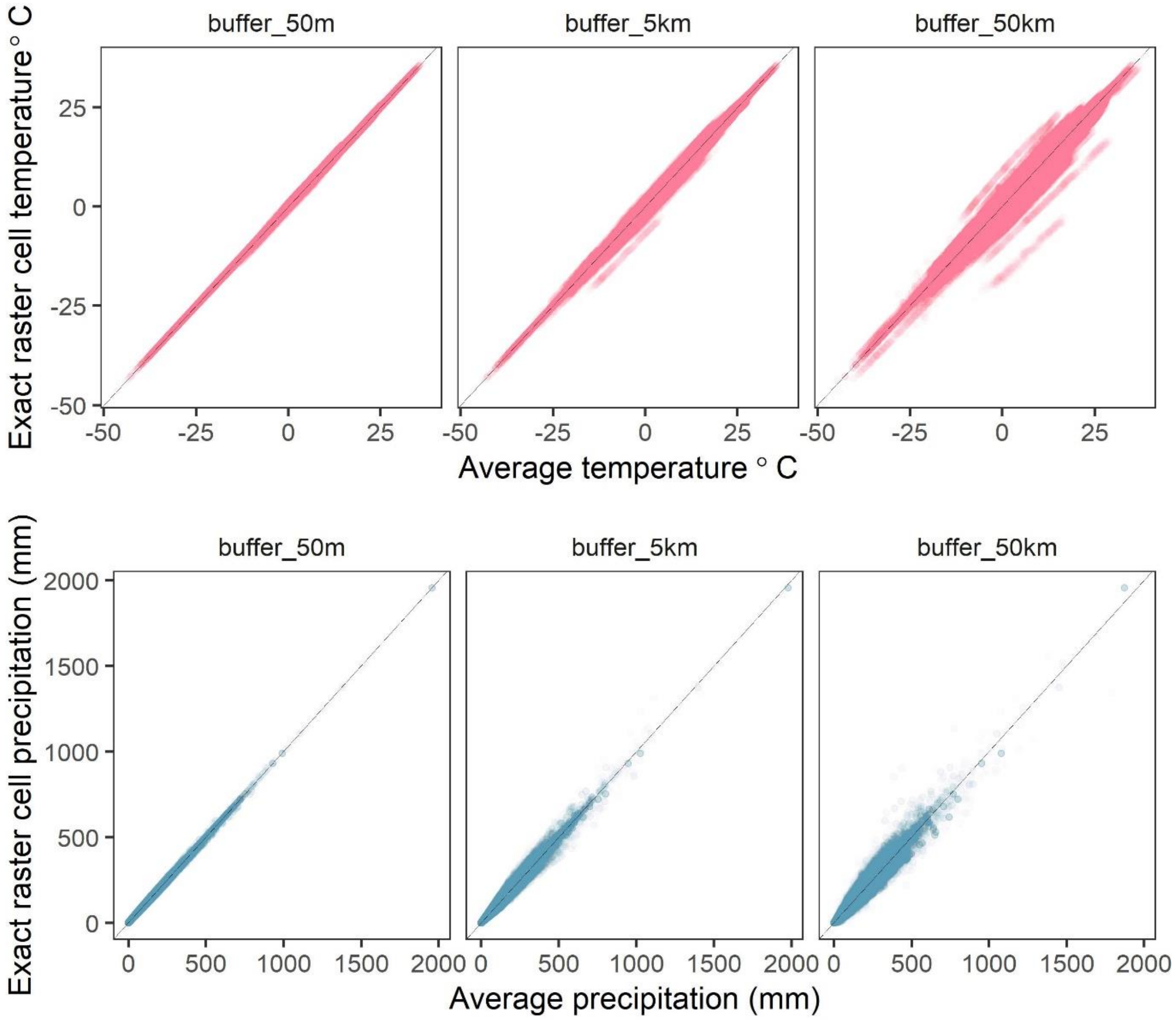


**Figure S1. Correlations between raster cell weather values for different buffer radii.** The average values of mean monthly temperature (top) and total precipitation (bottom) compared to exact raster cell values for buffer radii of 50m-50km (left-right) calculated from the CHELSA global gridded raster dataset. Buffered radii calculated using the *exactextractr* R package.

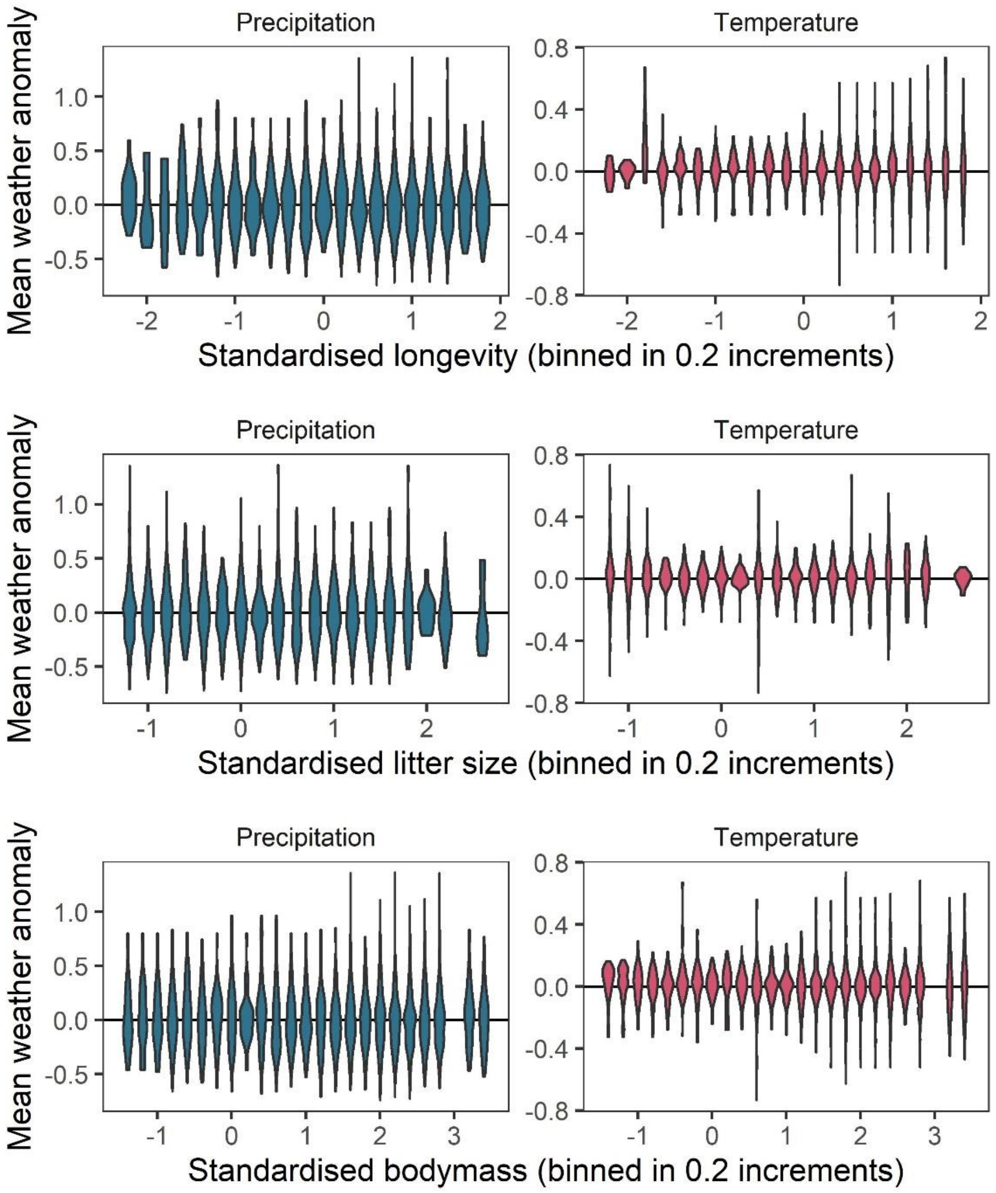


**Figure S2. Representation of the raw weather anomaly data across the observed ranges of life-history variables.** To verify whether a full range of weather anomalies was represented across the range of life-history variables observed, we examined the distributions of weather anomaly values observed for life-history variable bins of 0.2. These panels give violin distributions of weather anomalies for 0.2 increments of each of the three life-history variables (top-bottom) for temperature (right) and precipitation (left). These panels indicate a good spread of weather anomaly values observed across the life-history trait space.

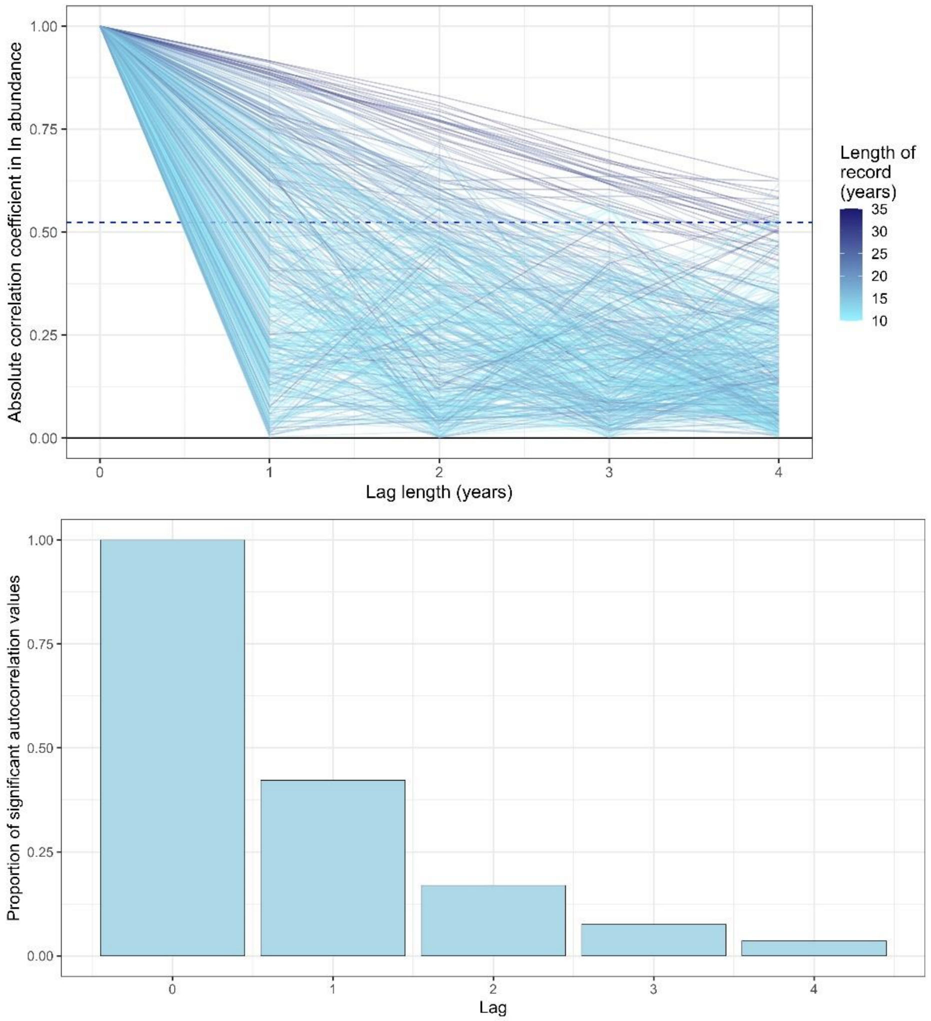


**Figure S3. Exploring temporal autocorrelation using autoregressive timeseries models of abundance with varying lag.** Top- the absolute autocorrelation coefficient for each lag length (in years) for each timeseries abundance record. Each line is a record from the mammal dataset used in the study, with lagged autocorrelation values up to a maximum lag of four years. The colour of the line indicates the total length of the record in years. The dashed line is the significance confidence level for the median timeseries length in the dataset (14 years). Bottom- the proportion of significant autocorrelation values across all records for each lag length (in years). Together, these figures show that there is good evidence for lag 1 autocorrelation (AR(1)) across all records, with a substantial proportion (>30%) of records displaying significant autocorrelation for AR(1). However, with greater lags, the degree of temporal autocorrelation decreases substantially, most probably due to the lack of sufficient annual observations to resolve autocorrelation with a greater lag.

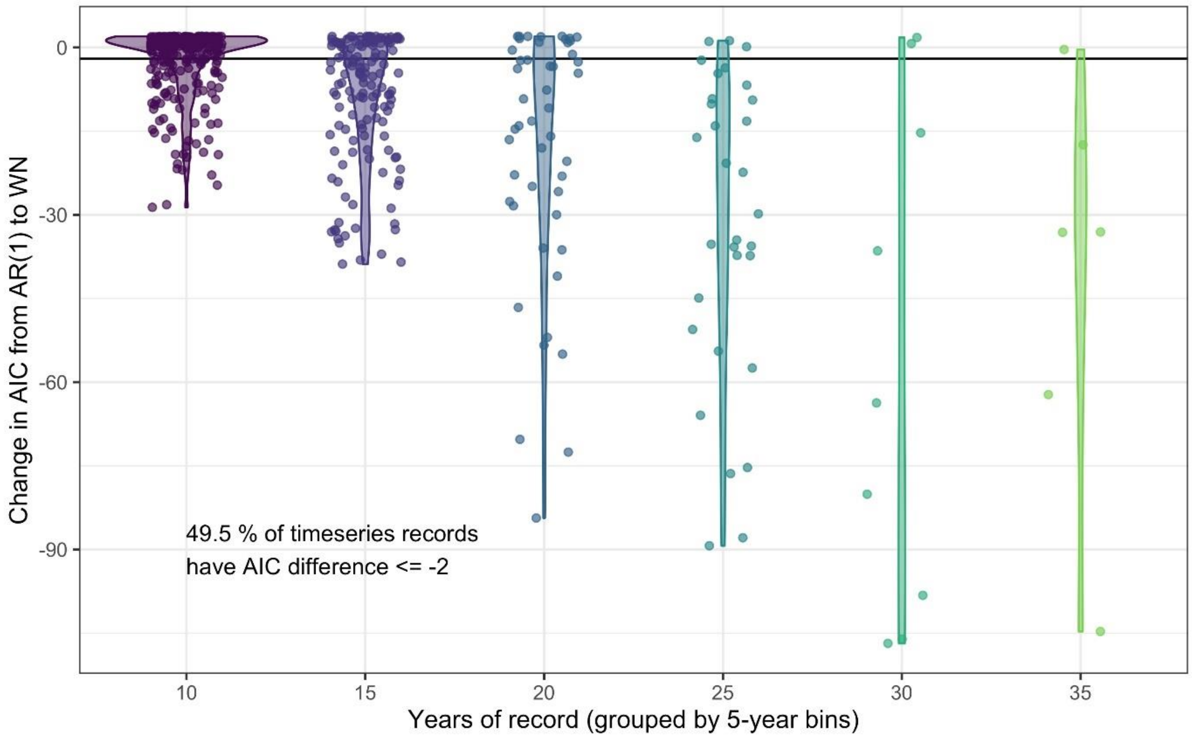


**Figure S4. Comparing the predictive performance of AR(1) time series models to white noise models for abundance records.** The change in AIC for timeseries models including an AR(1) temporal autocorrelation structure relative to the base model of white noise (WN). Each point gives the AIC difference for a single record, with the data grouped by the number of years (bins of five years) in the abundance record. Violins give the distribution of AIC differences for each record length bin. 49.5% of studies have an AIC difference <= -2 when comparing an AR(1) model to a white noise model, indicating support for including lag-1 temporal autocorrelation in models of abundance.

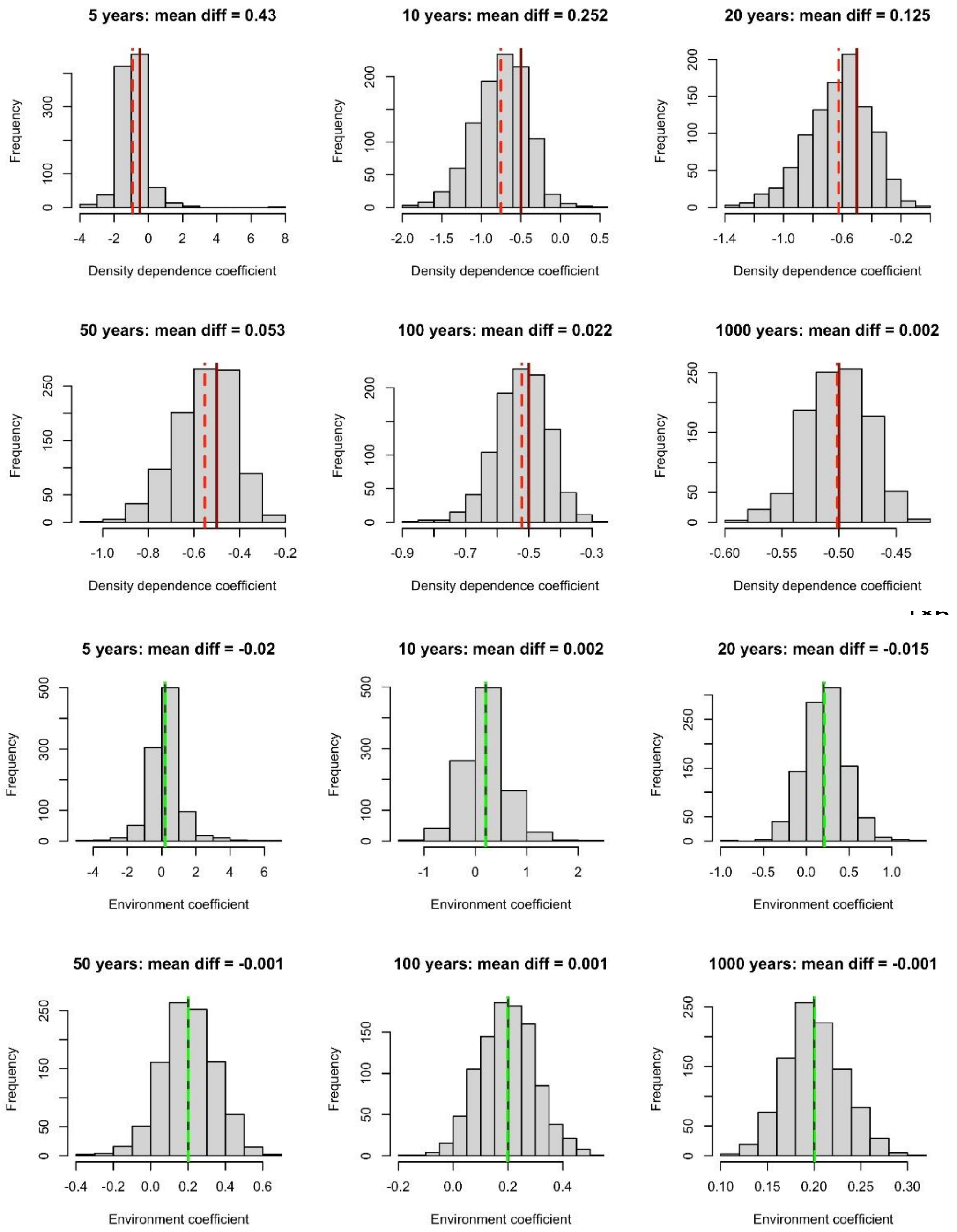


**Figure S5. Simulation results exploring the impact of incorporating density dependence on the estimation of environmental effects using linear models.** We built a time-series model simulation in which there was both lag-1 density dependence (coefficient = -0.5, solid red lines) and an environmental effect (coefficient = 0.2, solid green lines) and then retrofit linear models over 1000 simulations to explore whether we could estimate density dependence and environmental effects accurately for different time-series lengths. Top 6 panels (red lines) – the distribution of density dependence effects from retrofit linear models (mean = dashed red lines) over 1000 simulations for increasing timeseries lengths. Here, we did not accurately estimate density dependence effects for short timeseries. Bottom 6 panels (green lines) – the distribution of environmental effects from retrofit linear models (mean =

dashed green lines) over 1000 simulations for increasing timeseries lengths. Here, we accurately estimated environmental effects even for short time series. This suggests that accounting for temporal autocorrelation in abundance, we are able to retrieve accurate environmental effects (here weather effects).

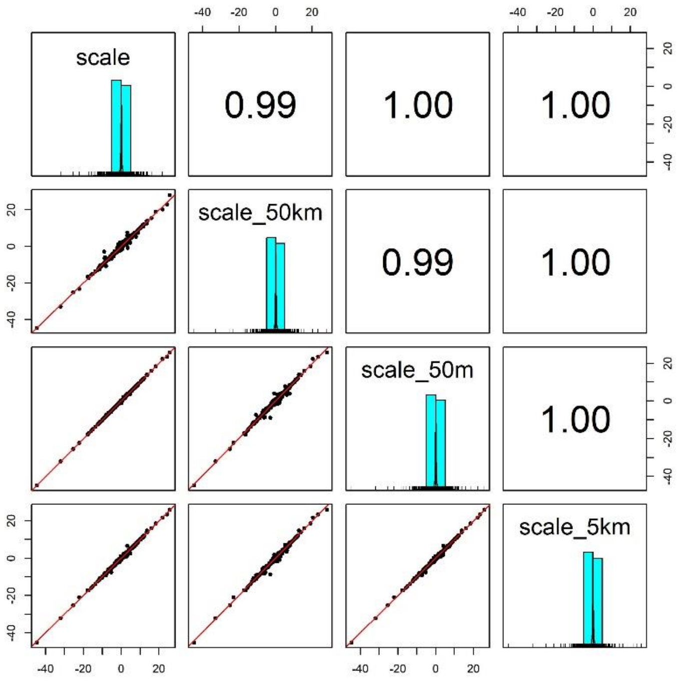


**Figure S6. Pairwise correlation coefficients for weather coefficients (temperature and precipitation) estimated over 4 buffered radii scales.** We estimated the effect of weather anomalies on population growth rate using additive models (GAMs) over 5 spatial scales for a buffered radius around each record's location- exact raster cell location (scale), 50m buffer (scale_50m), 5km buffer (scale_5km) and 50km buffer (scale_40km). Weather coefficients were near identical across spatial scales. The buffer radius of 5km was used in subsequent analyses.

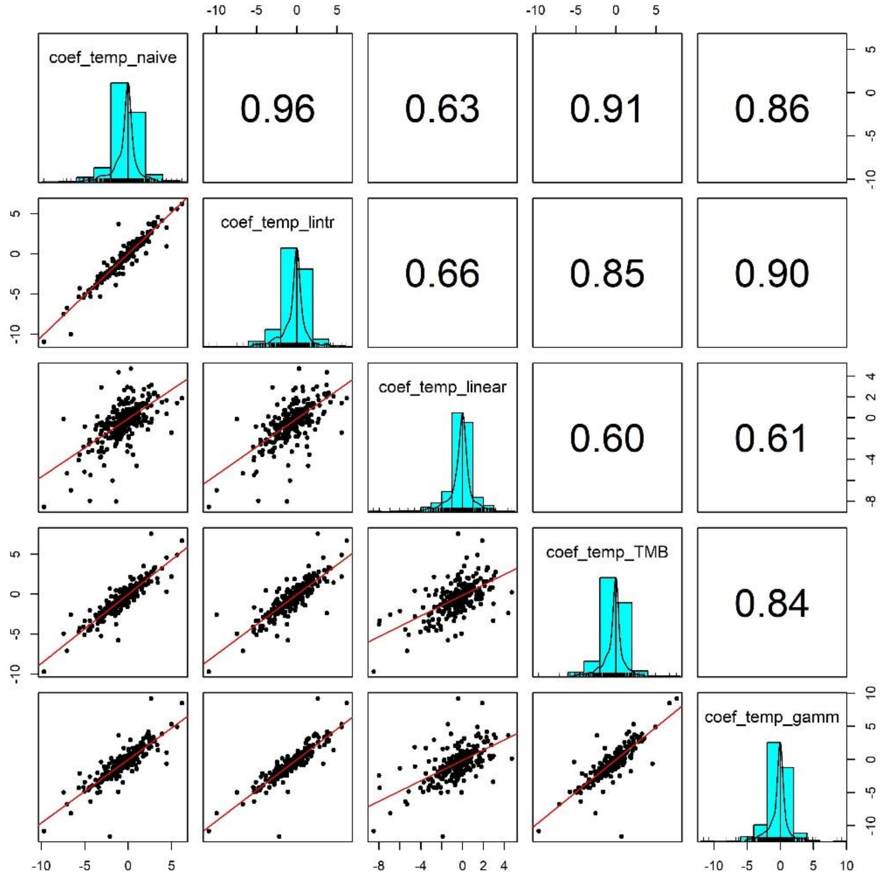


**Figure S7. Pairwise correlation coefficients for temperature effects estimated using 5 competing generalised linear models.** To validate our approach of using additive models to estimate weather effects, we compared the weather coefficients obtained from 5 models of differing complexity that accounted for (or excluded) temporal autocorrelation and temporal trends in population growth rates: 1) coef_temp_naive - simple linear regression excluding temporal trends or autocorrelation (R syntax population_growth_rate ~ weather_anomaly), 2) coef_temp_lintr – linear regression including a linear temporal trend in population growth rate but excluding autocorrelation (R syntax population_growth_rate ~ weather_anomaly + year), 3) coef_temp_linear – Linear regression including linear trend and an autoregressive term (R syntax population_growth_rate ~ weather_anomaly + year + abundance), 4) coef_temp_TMB - A glmmTMB model including an AR(1) autoregressive term for the observation year, 5) coef_temp_gamm – an additive model including a coarse smoothing spline for the temporal trend and an autoregressive term for year (see equation 2). Additive model coefficients were highly correlated with other estimates of weather effects (in support of Fig. S5)

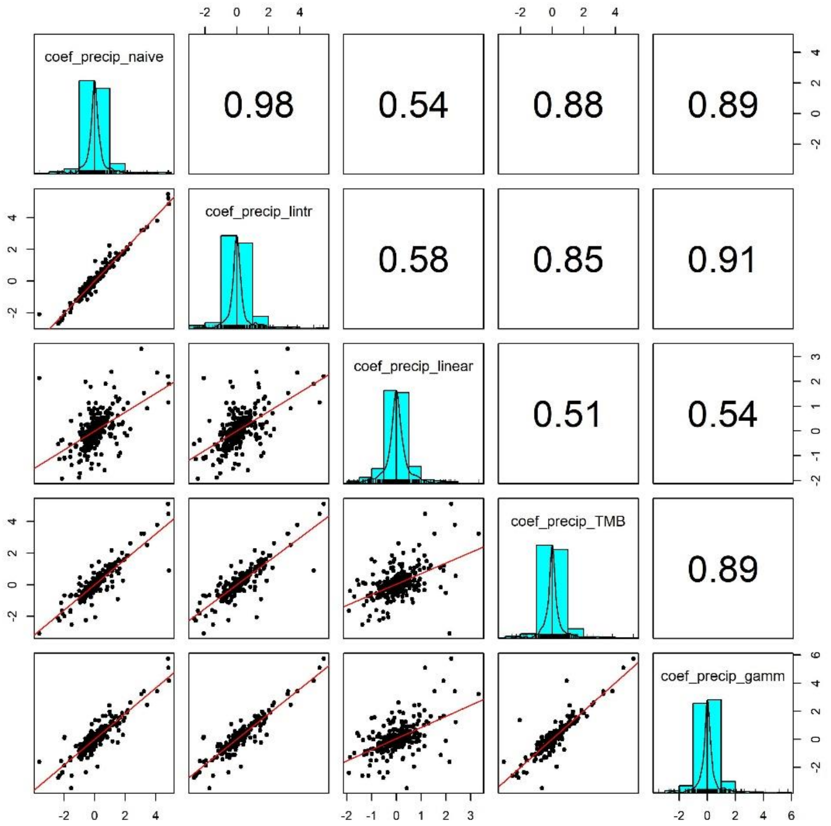


**Figure S8. Pairwise correlation coefficients for precipitation effects estimated using 5 competing generalised linear models.** To validate our approach of using additive models to estimate weather effects, we compared the weather coefficients obtained from 5 models of differing complexity that accounted for (or excluded) temporal autocorrelation and temporal trends in population growth rates (identical to Fig. S6 but using precip). Additive model coefficients for precipitation were highly correlated with other estimates of weather effects (in support of Fig. S5).

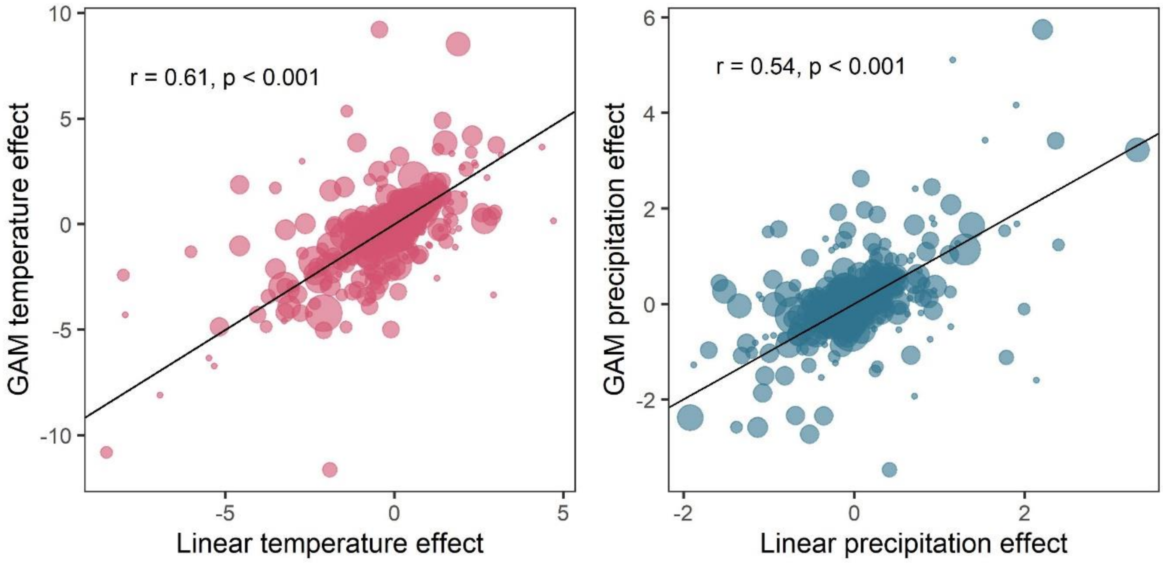


**Figure S9. Weather coefficients generated from linear regressions were positively correlated to those from additive models.** Comparison of weather coefficients generated from linear models including a linear trend term and an autoregressive term to additive models (equation 2). Highly significant positive correlation between linear and additive coefficients. This result is in support of the findings of the simulation presented in Fig. S5, which suggests that despite the method of accounting for density dependence (temporal autocorrelation), estimating annual environmental effects remains robust.

### S2 State-space approach

In addition to models fit using a generalised linear modelling approach, we also tested the validity of the final GAMM approach by calculating weather effect coefficients using state-space autoregressive time-series models that incorporated both process and observation error, which are often used as predictive models of time-series abundance data (e.g. (Daskalova et al., 2020)). Here, the state process of the population growth rate $r$ in year $t$ was a function of a linear effect of the weather variable on the abundance and random noise. State-space models were fit using the *rjags* and *jagsUI* packages in R (Kellner, 2021; Plummer, 2019) across 3 chains, which each had a total of 200,000 iterations, comprised of 100,000 burn-in iterations, 5000 adaptation iterations, and a thinning rate of 6. Across time-series records, there was a high fit-to sample, and the fit-to-sample was not influenced by the length of the time-series record (Fig. SX). We compared weather coefficients from state-space models with those obtained from GAMMs using Pearson's regression, and found highly significant correlations for both temperature and precipitation effects (Fig. SX).

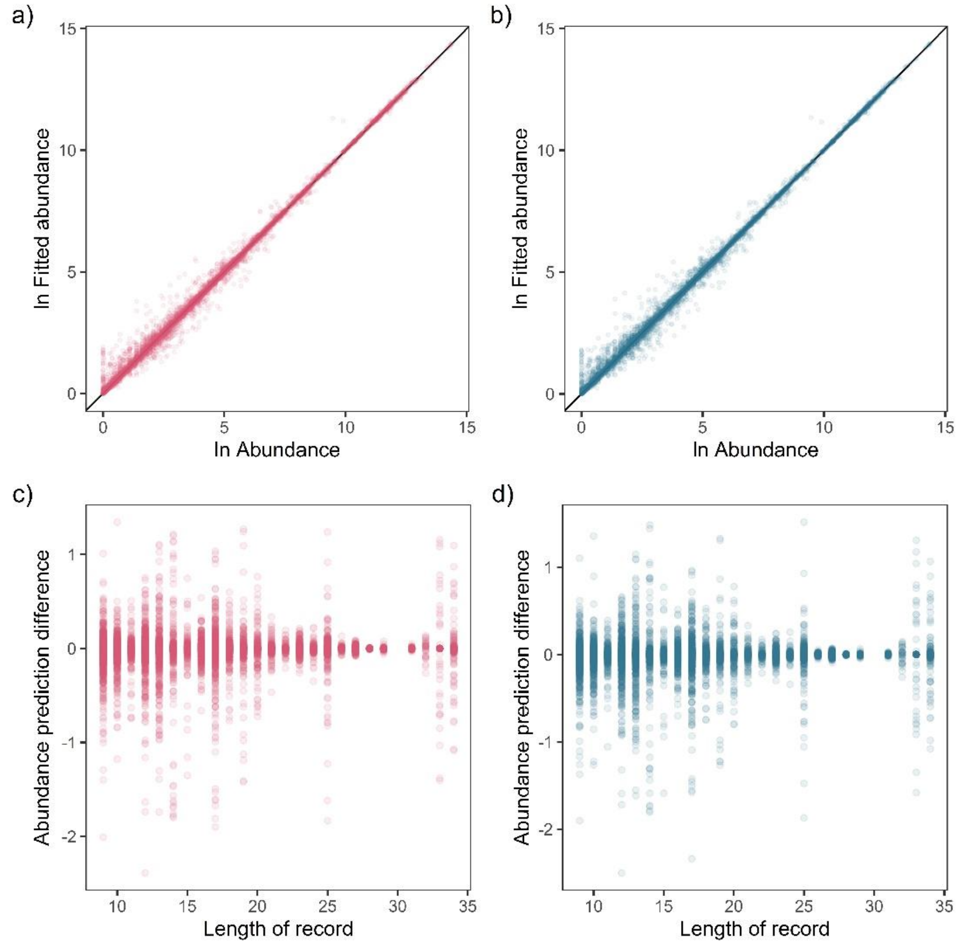


**Figure S10. Fit-to-sample estimates for state-space models on mammal abundance time-series.** Top) Fitted vs. observed annual ln abundance values for temperature (a) and precipitation (b) effects across all observations of 474 (non-NA in precipitation anomaly) records, with solid black line giving the 1-to-1 line. Bottom) The relationship between the difference in observed and fitted ln abundance with respect to the length of the time-series record for temperature (c) and precipitation (d).

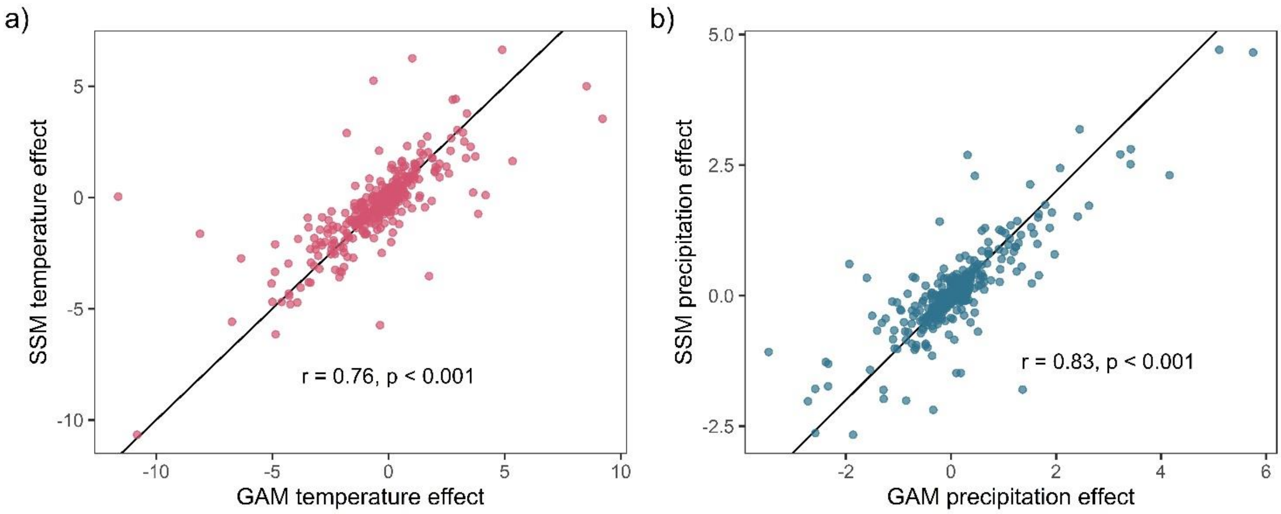


**Figure S11. Weather coefficients generated from state-space models (SSMs) were highly correlated to those from GAM models.** Highly significant positive Pearson's correlation between state-space and additive coefficients for both temperature (a) and precipitation (b). This result supports the validity of the use of GAM models to estimate weather effects in the current study.

### S3 Prior predictive simulation

We used conservative, regularising priors following McElreath 2020, which gave predictions within the parameter space observed in the raw data. This was achieved through prior predictive simulation (PPS; Fig. S12-S15). Here, we compare the estimates and predictions of priors to the limits of observed data and expected patterns to inform the priors. In addition to prior choices made in this section, we further tuned the priors during the model selection to improve the efficiency/accuracy of Markov chains. Choosing conservative regularising priors also reflected the large number of parameters in phylogenetically or spatially controlled models. Specifically, we performed PPS for the global intercept term of consistent weather effects, the $\beta$ terms relating to differences in weather effects (i.e. biome effects), $\beta$ terms for linear life-history effects in equivalent log-normal models, and mixed-effects variance terms for species variance and phylogenetic covariance.

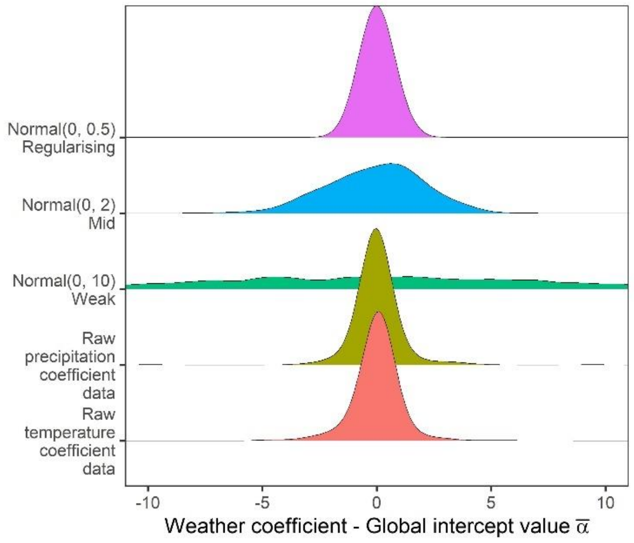


**Figure S12. Prior predictive simulation for global intercept terms.** The global intercept i.e. consistent pattern of weather effects across records was modelled using a normal prior. Density distributions for the weather coefficients observed in the raw data, and weather coefficients under 3 normal priors, weak (mean = 0, sd = 10), medium (mean = 0, sd = 2), regularising (mean = 0, sd = 0.5). Here, the regularising prior gives likely global intercepts within the range of the observed coefficients.

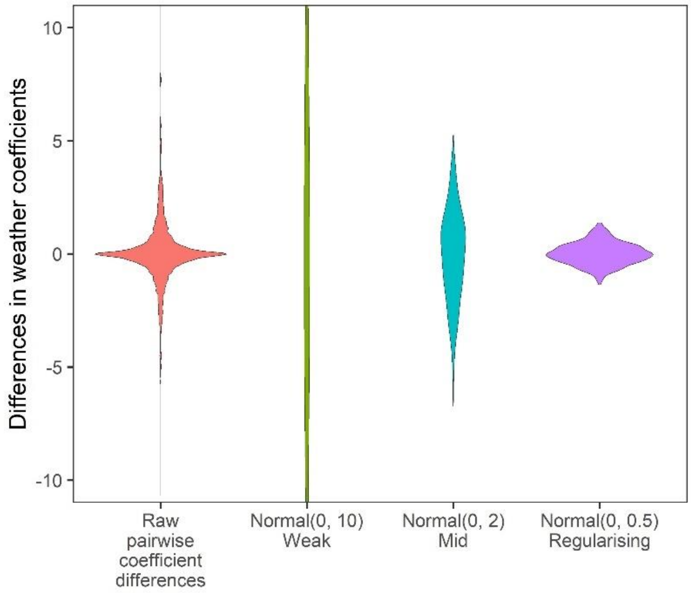


**Figure S13. Prior predictive simulation for $\beta$ terms giving differences in weather coefficients.** Density distributions for all pairwise differences in the weather coefficients observed in the raw data, and differences in weather coefficients under 3 normal priors, weak (mean = 0, sd = 10), medium (mean = 0, sd = 2), regularising (mean = 0, sd = 0.5). The regularising prior gives difference values within the range of the observed coefficient differences.

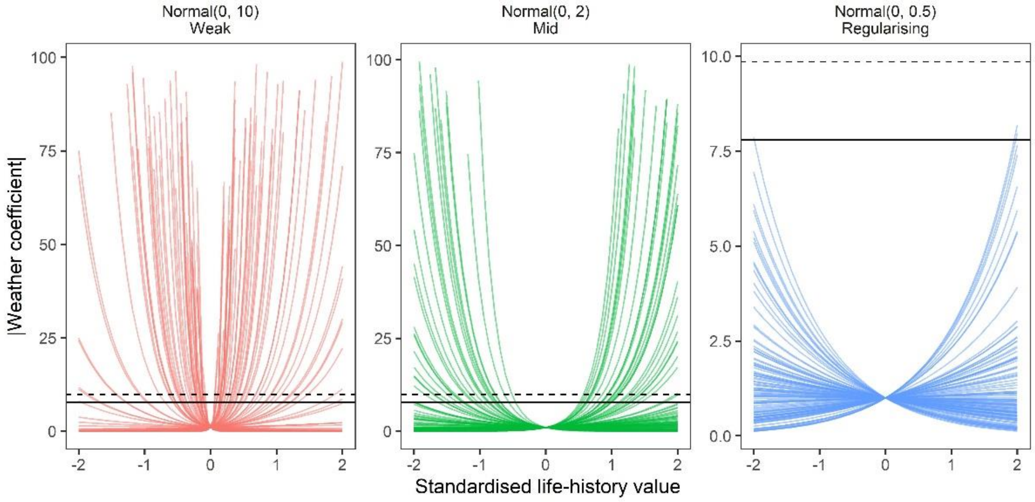


**Figure S14. Prior predictive simulation for $\beta$ terms giving life-history effects on weather coefficients.** Prior predictions of the effect of simulated scaled life-history values (-2 to 2) on absolute weather coefficients as using log-normal models. Lines are individual simulations under normal prior distributions for the linear life-history effect, which was back-transformed using the exponential to give absolute weather coefficients. Panels give the prior simulations of life-history on weather coefficients under 3 normal priors, weak (mean = 0, sd = 10), medium (mean = 0, sd = 2), regularising (mean = 0, sd = 0.5). solid and dashed horizontal lines give the maximum observed absolute coefficients for temperature and precipitation, respectively. The regularising prior gives plausible predictions that do not regularly exceed the maximum and minimum effects observed.

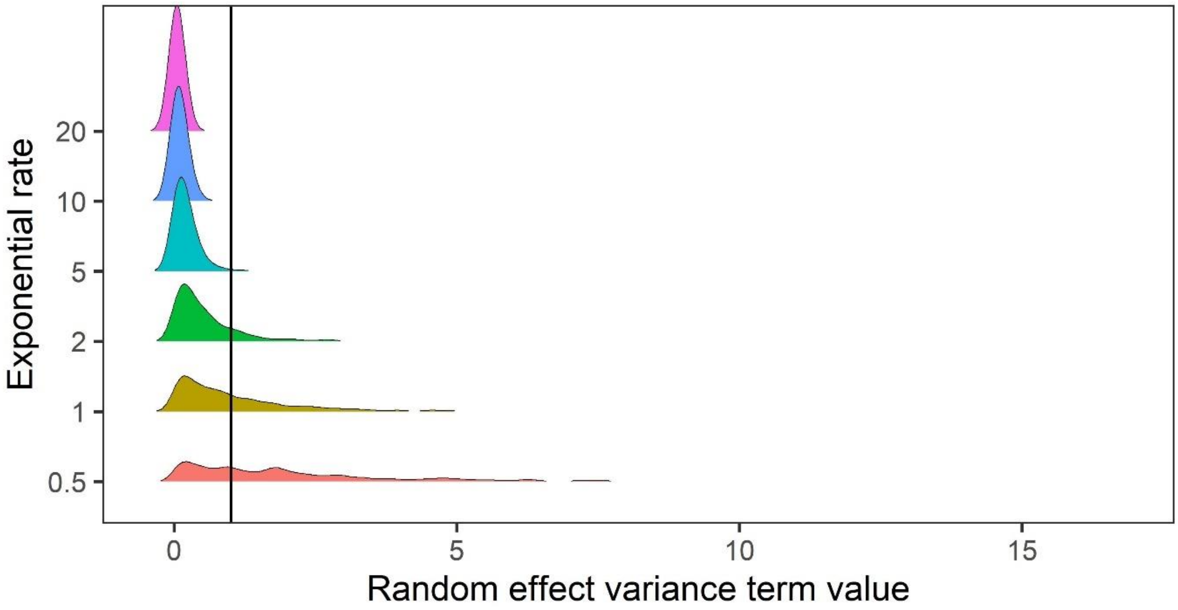


**Figure S15. Prior predictive simulation for standard deviation terms relating to random effects of variance.** The random effects of species-level variance and phylogenetic covariance were modelled using exponential priors, which are suitable for variance terms because they are non-zero distributions that can flexibly capture higher variances. Here, we explored the density distributions of exponential priors with 6 exponential rate parameters (0.5-20). In this case, for phylogenetic and species level variance we do not expect values exceeding a variance term of 1 (solid black line). Regularising priors with rate parameters >= 5 gave conservative estimates of random effect variances within the constraints of the variance terms in the meta-regression models.

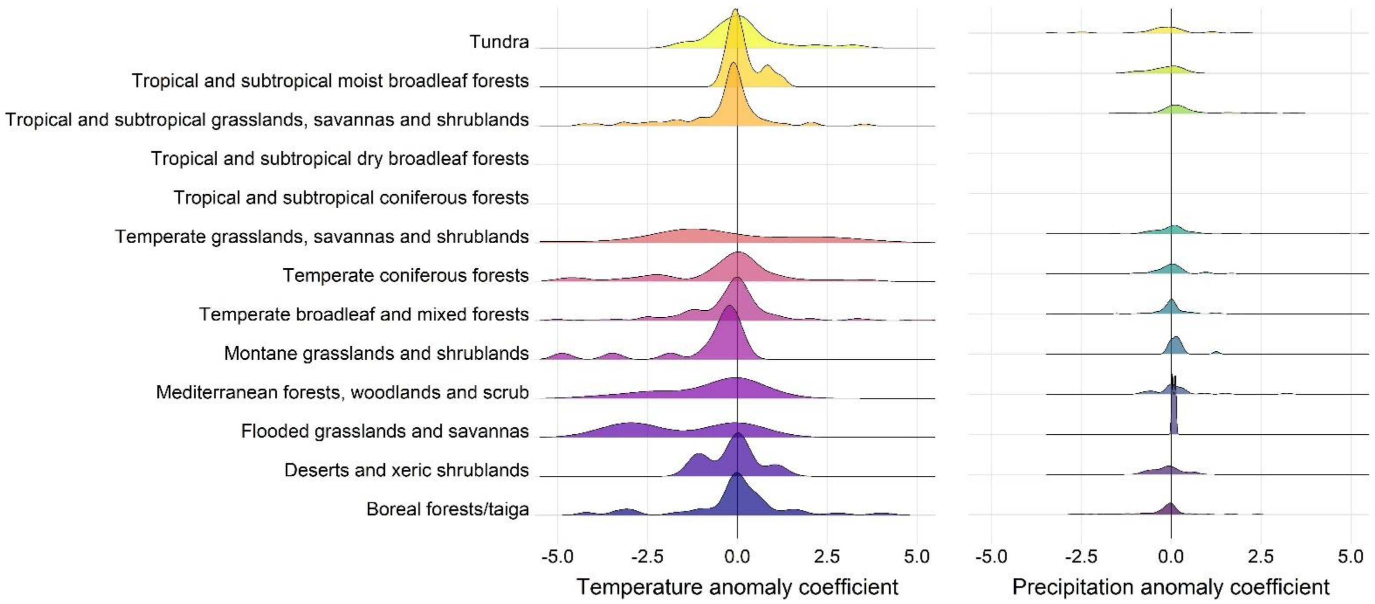


**Figure S16. Density distributions of temperature (left) and precipitation (right) coefficients on abundance change with respect to biome of the record location in the terrestrial mammals.** Only coefficients between -0.5-0.5 are displayed for visual purposes.

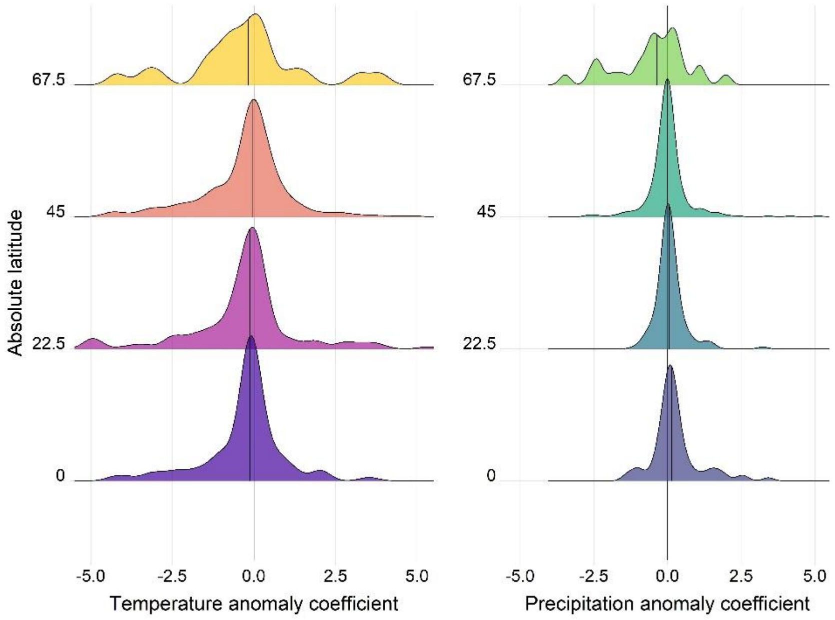


**Figure S17. Density distributions of temperature (left) and precipitation (right) coefficients on abundance change with respect to the absolute latitude of the record location in the terrestrial mammals.** Records were grouped based on their absolute latitude in categories of 22.5° e.g. 0 category indicates records found at absolute latitudes of 0-22.5°.

**Table S1. Model selection results from Gamma models exploring the impact of life-history traits on absolute responses to temperature in the terrestrial mammals.** Leave One Out (LOO) cross validation results for 10 candidate models including life-history traits relative to a base model excluding life-history traits. elpd = expected log pointwise predictive density.

| Model | predictors | LOO elpd | LOO elpd error | elpd difference | elpd error difference | LOO information criterion |
|---|---|---|---|---|---|---|
| temp_longevity | longevity | -179.10 | 27.29 | 0.00 | 0.00 | 358.21 |
| temp_lh_uni | longevity + bodymass + litter | -180.00 | 27.65 | 0.90 | -0.36 | 360.01 |
| temp_litbod_simple | litter + bodymass | -180.49 | 27.42 | 1.38 | -0.13 | 360.98 |
| temp_litter | litter | -180.56 | 28.00 | 1.46 | -0.71 | 361.12 |
| temp_litbod | litter*bodymass | -180.61 | 27.38 | 1.51 | -0.10 | 361.23 |
| temp_lonbod_simple | longevity + bodymass | -180.82 | 27.11 | 1.72 | 0.18 | 361.65 |
| temp_lonbod | longevity*bodymass | -181.00 | 27.03 | 1.90 | 0.26 | 362.00 |
| temp_lh | longevity + bodymass + litter + longevity:bodymass + litter:bodymass + litter:longevity | -181.57 | 27.68 | 2.46 | -0.39 | 363.13 |
| temp_bodymass | bodymass | -183.52 | 26.99 | 4.41 | 0.29 | 367.03 |
| temp_base | | -184.54 | 27.21 | 5.44 | 0.07 | 369.09 |

**Table S2. Model selection results from Gamma models exploring the impact of life-history traits on absolute responses to precipitation in the terrestrial mammals.** Leave One Out (LOO) cross validation results for 10 candidate models including life-history traits relative to a base model excluding life-history traits. elpd = expected log pointwise predictive density.

| Model | predictors | LOO elpd | LOO elpd error | elpd difference | elpd error difference | LOO information criterion |
|---|---|---|---|---|---|---|
| precip_longevity | longevity | -108.26 | 29.79 | 0.00 | 0.00 | 216.53 |
| precip_lonbod_simple | longevity + bodymass | -108.50 | 29.81 | 0.24 | -0.03 | 217.00 |
| precip_litter | litter | -108.51 | 29.80 | 0.24 | -0.01 | 217.01 |
| precip_litbod_simple | litter + bodymass | -108.54 | 29.70 | 0.27 | 0.09 | 217.07 |
| precip_lh_uni | longevity + bodymass + litter | -108.61 | 29.75 | 0.35 | 0.03 | 217.22 |
| precip_lonbod | longevity*bodymass | -108.93 | 29.84 | 0.67 | -0.05 | 217.86 |
| precip_bodymass | bodymass | -109.16 | 29.72 | 0.89 | 0.07 | 218.31 |
| precip_base | | -109.30 | 29.84 | 1.03 | -0.05 | 218.59 |
| precip_lh | longevity + bodymass + litter + longevity:bodymass + litter:bodymass + litter:longevity | -110.08 | 29.96 | 1.81 | -0.17 | 220.16 |
| precip_litbod | litter*bodymass | -110.14 | 29.87 | 1.87 | -0.08 | 220.27 |

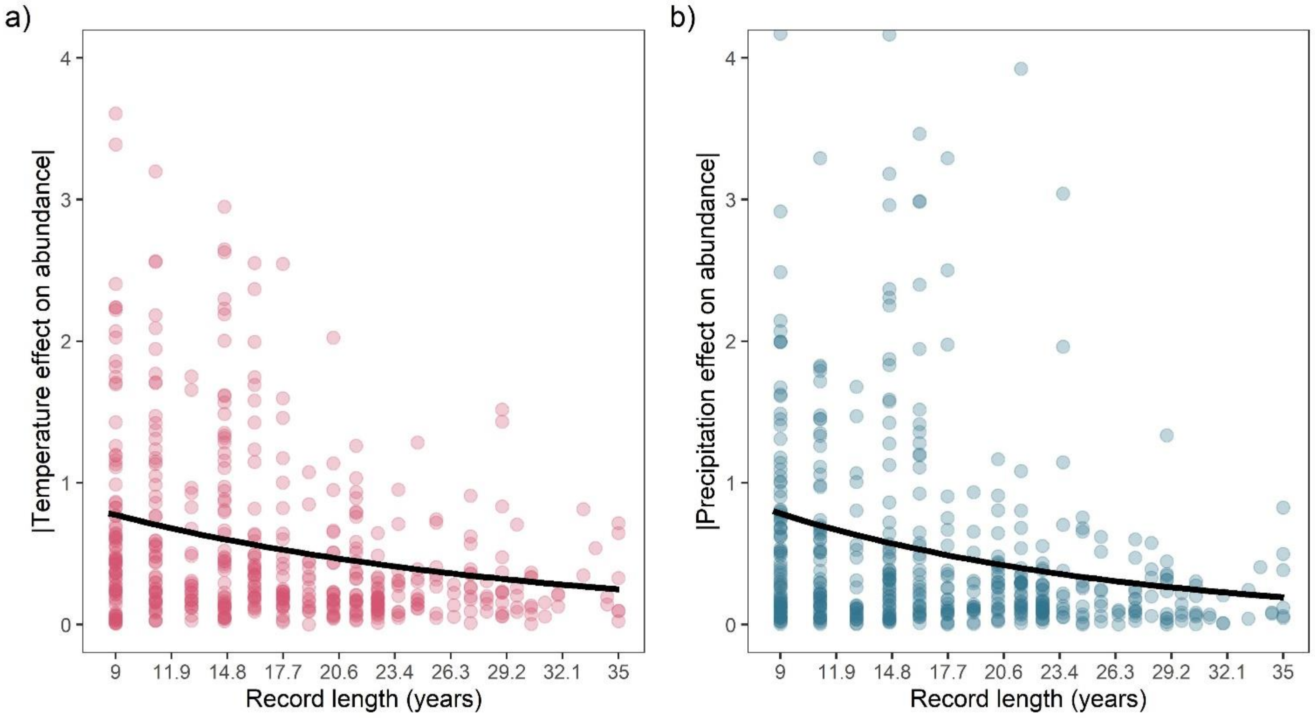


**Figure S18. Posterior predictions for the influence of the record length on absolute temperature (a) and precipitation (b) on abundance changes in the terrestrial mammals.** Points give the absolute weather effect for each record (N = 486). Only absolute weather effects <4 are displayed on the figure. Black lines are posterior means from the best predictive Gamma model including life-history effects.

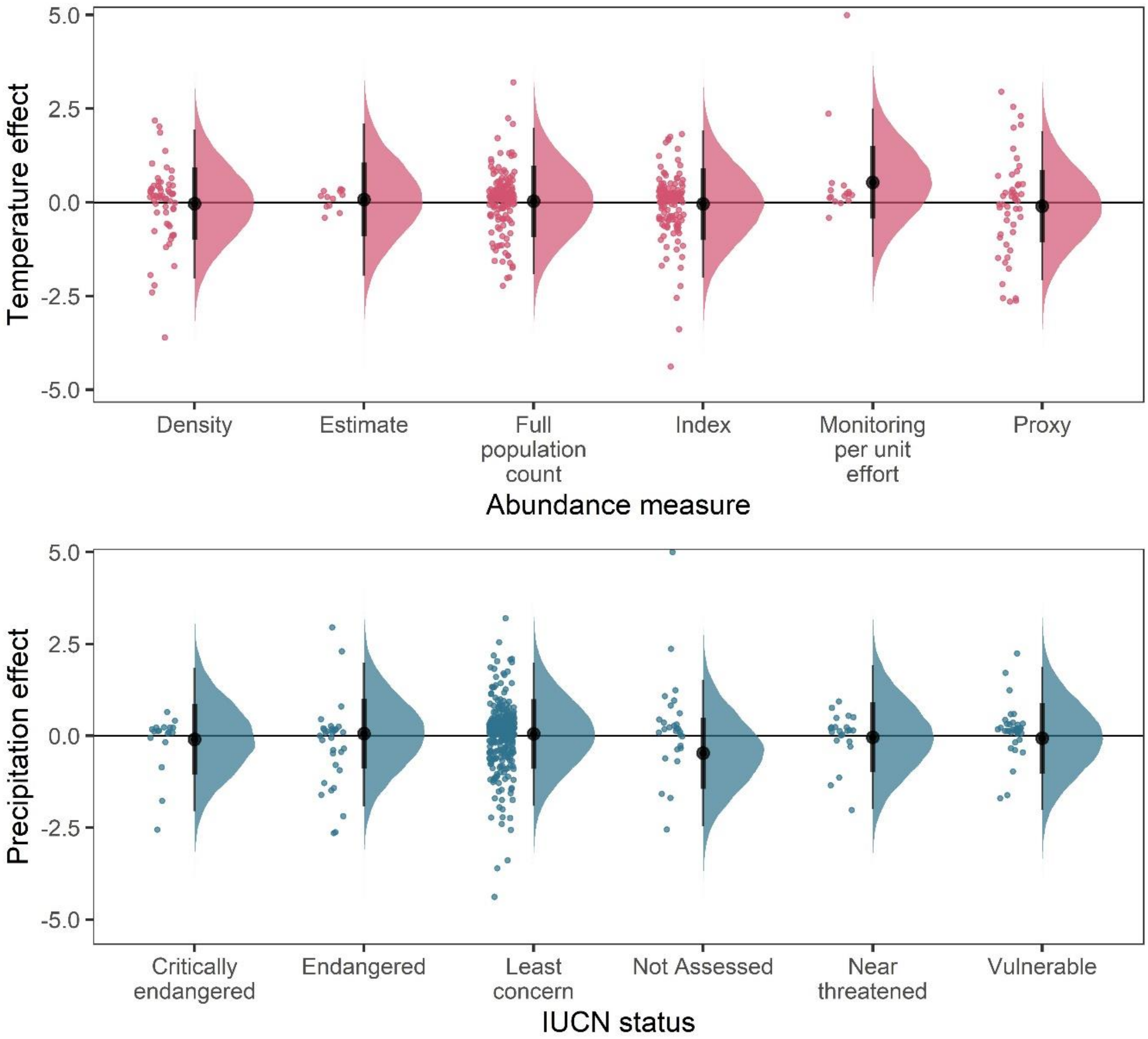


**Figure S19. Posterior predictions for a Gaussian meta-regression model of temperature coefficients with the method of estimating abundance (top) and temperature coefficients with IUCN red-list status**. To explore the potential impact of varying data-sources and reliability of the abundance data, we performed a further model selection to explore how the abundance measure type and IUCN red-list status influence temperature and precipitation responses (as in equation 3). For temperature coefficients the model including abundance measure type had a higher predictive performance than the base model (Δelpd = 3.13). For precipitation coefficients the model including IUCN red-list status had a higher predictive performance than the base model (Δelpd = 0.13). However, posterior predictions revealed that these predictive differences were not substantive, with posterior distributions centred on 0 for most abundance measure types (top) and IUCN statuses (bottom). The predictive differences are most likely a result of coefficients observed in the Monitoring per unit effort measure type (top) and the Not Assessed status (bottom), both of which have low sample size in the terrestrial mammals.

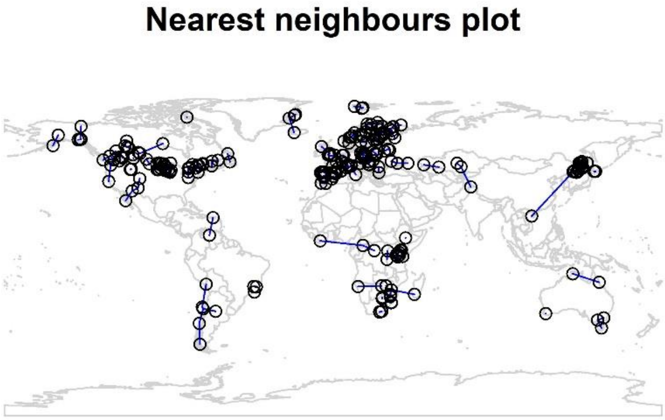


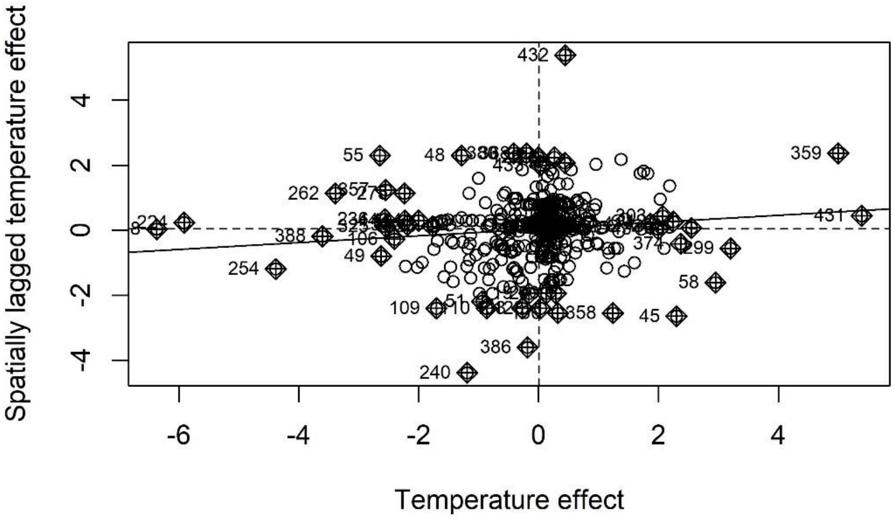


**Figure S19. Nearest neighbour plot for abundance record localities and Moran's I plot for temperature coefficients.** We explored spatial autocorrelation in the coefficients of the GAM models (equation 2) using Moran's I calculations with a nearest neighbours approach. Top – nearest neighbours plot for each abundance record in the study. We found low magnitude Moran's I for both temperature ($I = 0.11$, $p = 0.03$) and precipitation ($I = 0.05$, $p > 0.05$), but a significant Moran's I for temperature. Bottom – Moran's I plot for temperature effects indicates weak correlation between temperature effects and spatially lagged temperature effects, but the Moran's I plot indicates this is due to a small number of studies with high spatially lagged values.

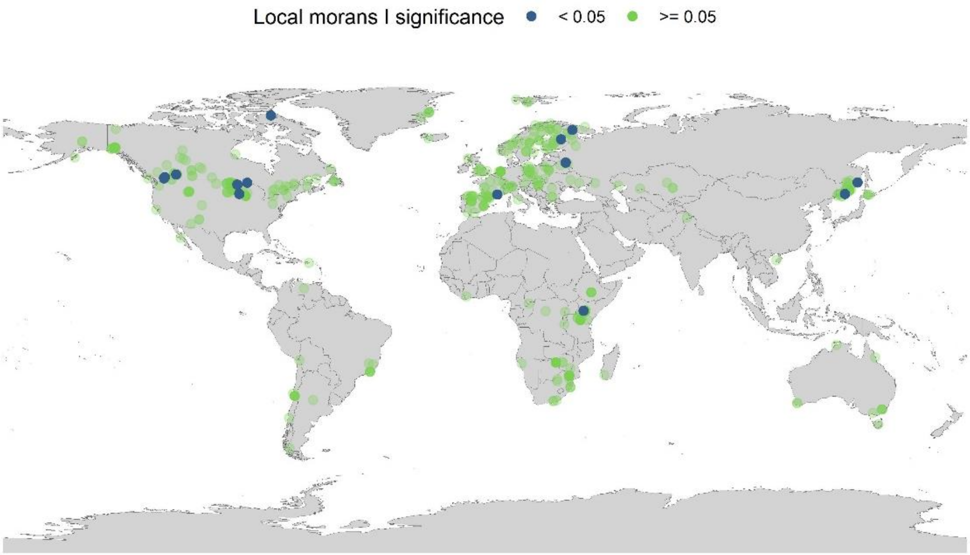


**Figure S20. Local Moran's I significance for temperature coefficients**. In addition to the global Moran's I analysis for temperature, we ran a local Moran's I for the temperature coefficients. Each point on the map gives the local Moran's I significance rating (95% level) and the spatial location of all records in the study. The local Moran's I indicates that a small number of spatially autocorrelated points is dictating general spatial autocorrelation patterns in the abundance records.

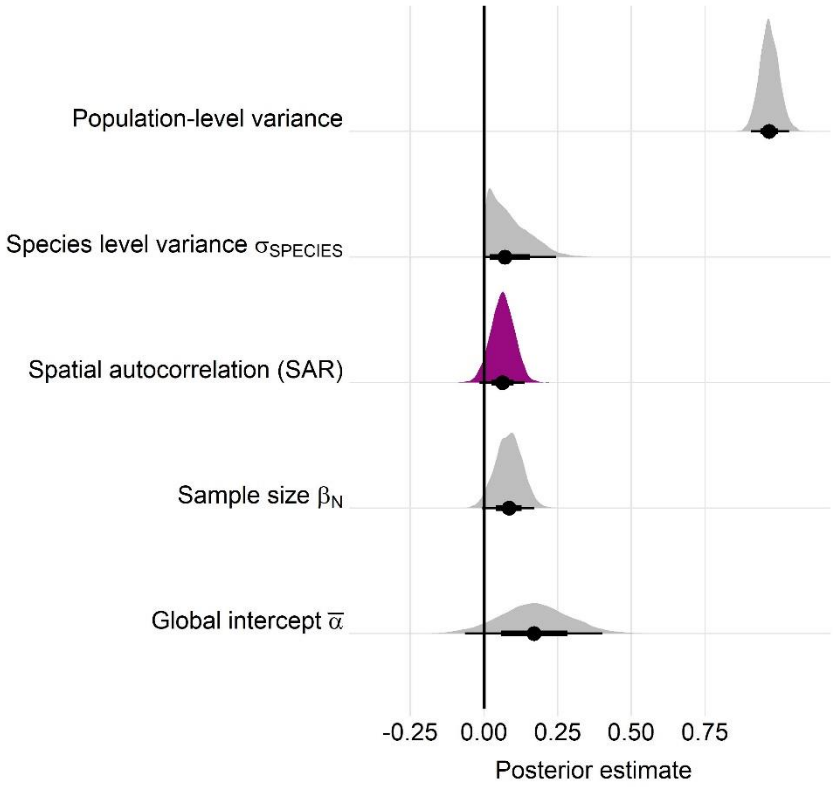


**Figure S21. Posterior estimates for a Gaussian meta-regression model of temperature coefficients with an explicit spatially autocorrelated term across the terrestrial mammals**. Posterior density distributions for each key model parameter (as in equation 2) given with half eye plots, where the point indicates the posterior average, and the bar is calculated using a cumulative distribution function. Purple density indicates the posterior estimate for the spatially autocorrelated term, whose posterior distribution overlapped with zero. Furthermore, leave-one-out cross validation indicated that the base model excluding spatial autocorrelation had a higher predictive performance than the model including the spatial term.

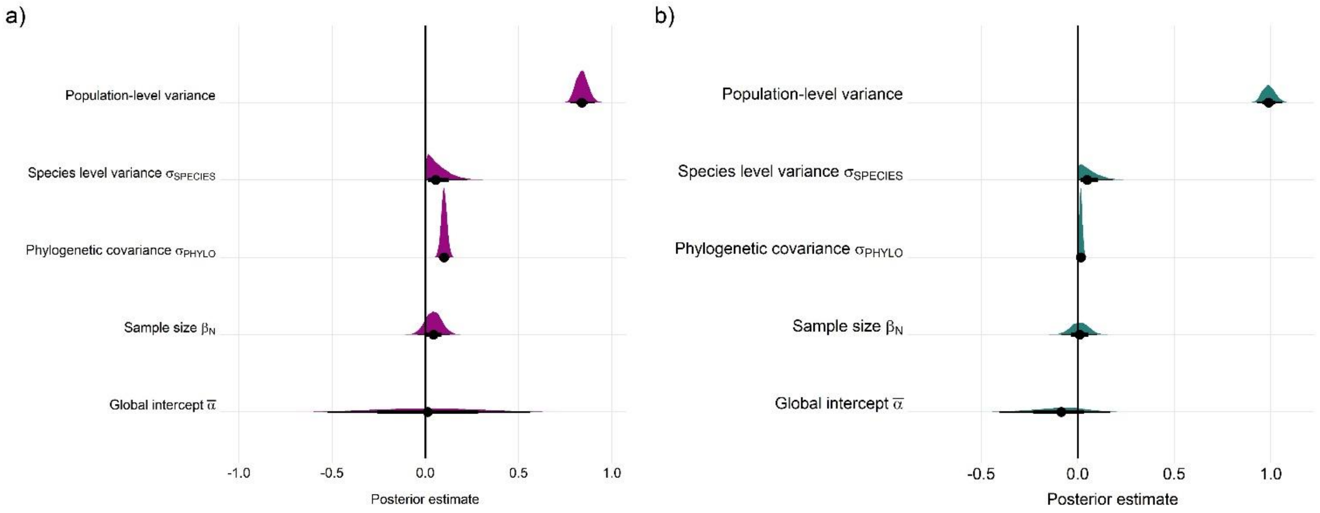


**Figure S22. Posterior estimates for Gaussian meta-regression models for temperature (a) and precipitation (b) variance across the terrestrial mammals**. Posterior density distributions for each key model parameter (as in equation 2) given with half eye plots, where the point indicates the posterior average, and the bar is calculated using a cumulative distribution function.
